# The *Saccharomyces cerevisiae* Hrq1 and Pif1 DNA helicases synergistically modulate telomerase activity *in vitro*

**DOI:** 10.1101/327320

**Authors:** David G. Nickens, Cody M. Rogers, Matthew L. Bochman

## Abstract

Telomere length homeostasis is vital to maintaining genomic stability and is regulated by multiple factors, including telomerase activity and DNA helicases. The *Saccharomyces cerevisiae* Pif1 helicase was the first discovered catalytic inhibitor of telomerase, but recent experimental evidence suggests that Hrq1, the yeast homolog of the disease-linked human RecQ-like helicase 4 (RECQL4), plays a similar role via an undefined mechanism. Using yeast extracts enriched for telomerase activity and an *in vitro* primer extension assay, here we determined the effects of recombinant wild-type and inactive Hrq1 and Pif1 on total telomerase activity and telomerase processivity. We found that titrations of these helicases alone have equal-but-opposite biphasic effects on telomerase, with Hrq1 stimulating activity at high concentrations. When the helicases were combined in reactions, however, they synergistically inhibited or stimulated telomerase activity depending on which helicase was catalytically active. These results suggest that Hrq1 and Pif1 interact and that their concerted activities ensure proper telomere length homeostasis *in vivo*. We propose a model in which Hrq1 and Pif1 cooperatively contribute to telomere length homeostasis in yeast.

---

Telomerase is a ribonucleoprotein complex (1) that functions to overcome the end replication problem (2) encountered by linear eukaryotic chromosomes. Although the telomerase holoenzyme differs between organisms (or has been entirely replaced by retrotransposons, as in *Drosophila* (3)), all versions studied to date contain two conserved subunits: a reverse transcriptase and a telomerase RNA that is used as the template for the reverse transcriptase (1). In humans, these subunits are termed TERT and *TER*, respectively, but in yeast, they are referred to as Est2 and *TLC1*. While telomerase plays a critical role in genome duplication, it is also abnormally activated in most cancers, allowing cancer cells to overcome the normal cell division limit (*i.e*., the Hayflick limit) of a somatic cell before it senesces (4). As such, telomerase is tightly regulated to prevent rampant immortalization of cells and to maintain telomere length homeostasis.

Telomerase regulation can occur by many methods. For instance, the transcription of human TERT can be altered genetically and epigenetically in cancers, leading to upregulation of telomerase activity (5). Similarly, the transcription of genes encoding telomerase components and telomerase activity itself can be up- or downregulated by various dietary compounds (6). Post-translational modifications of TERT, such as phosphorylation (7) and ubiquitination (8), also affect telomerase activity. However, there is a rich and growing body of literature describing the effects of DNA helicases on telomerase (reviewed in (9)). Many of these helicases, including the RecQ family members BLM and WRN and the iron-sulfur cluster family helicases RTEL1 and FANCJ, are thought to unwind DNA secondary structures such as G-quadruplexes and t-loops that are formed at telomeres.

In contrast, the *Saccharomyces cerevisiae Pif1* helicase is hypothesized to function by a different mechanism. In the absence of Pif1, yeast telomeres lengthen over successive generations, suggesting that Pif1 functions as a telomerase inhibitor (10). Biochemical work demonstrates that Pif1 likely evicts telomerase from telomeres by unwinding the *TLC1* RNA-telomeric DNA hybrid that must be formed to enable Est2 reverse transcription (11,12). As Pif1 preferentially localizes to the longest telomeres *in vivo* (13), this effectively titrates telomerase away from long telomeres toward the short telomeres that are most in need of lengthening, resulting in a stable homeostatic telomere length.

Recently, we discovered that a second yeast helicase, Hrq1, functions in telomere length homeostasis (14). Although deletion of *HRQ1* has no effect on bulk telomere length, the absence of Hrq1 in the *pif1-m2* genetic background (which lacks the nuclear isoform of Pif1) leads to hyper-lengthening of telomeres relative to *pif1-m2* cells alone. Further, using gross-chromosomal rearrangement (GCR) assays, we found that GCR events are preferentially healed by telomere addition in *hrq1Δ* cells relative to predominantly recombination-mediated repair in wild-type. Hrq1 also promotes the formation of type I survivors in telomerase-null (*tlc1Δ*) cells. As Hrq1 can be localized to telomeres by chromatin-immunoprecipitation, and telomeric repeat sequence DNA (TG_1-3_) is a preferred *in vitro* substrate for Hrq1 (15), it likely affects telomerase activity directly.

Despite these parallels with Pif1 inhibition of telomerase, we hypothesize that the mechanisms by which the two helicases exert this effect are different. For instance, the ATPase-null allele of *PIF1* (*pif1-K264A*) phenocopies both *pif1Δ* and *pif1-m2*, indicating that catalytic activity by Pif1 is necessary for telomerase inhibition (11,16,17). In contrast, in the GCR assay, cells expressing the ATPase-null allele of *HRQ1* (*hrq1-K318A*) display very low, nearly wild-type levels of telomere addition at DNA double-strand breaks (DSBs) compared to the ~80% of telomere additions recovered in *hrq1Δ* cells (14). This would suggest that Hrq1 has a structural rather than catalytic role in DSB repair and telomere maintenance.

These data are particularly evocative in light of the fact that the human homolog of Hrq1, RECQL4 (or RECQ4), has an important but largely unknown role in telomere maintenance (18,19), and defects in this process could underlie the genomic instability characteristic of RECQL4-related diseases (20). Thus, studying the mechanism(s) by which Hrq1 modulates telomerase activity in yeast may shed light on the role of RECQL4 in telomere homeostasis. Here, we used an *in vitro* primer extension assay (21) to characterize the activity of telomerase in the presence of wild-type and inactive Hrq1, Pif1, and combinations of both. We found that recombinant Hrq1 or Hrq1-K318A had modest effects on telomerase activity alone, but addition of both Hrq1 and Pif1 to the assay yielded synergistic inhibition or activation of telomerase activity depending on the catalytic activity of the helicases. We present a model in which Hrq1 and Pif1 cooperate to contribute to telomere length homeostasis in yeast.

## RESULTS

### Hrq1 affects telomerase activity by a different mechanism than Pif1

As described above, genetic and biochemical evidence suggests that Hrq1 and Pif1 both play roles in telomere maintenance, but it is likely that Hrq1 affects telomerase activity by a mechanism distinct from that of Pif1. Indeed, the *in vitro* helicase activity of Hrq1 is decreased approximately fourfold when unwinding a fork substrate with an RNA-DNA hybrid duplex region relative to a DNA-DNA duplex (Fig. 1A). In contrast, Pif1 helicase activity is stimulated by RNA-DNA hybrids (12,22).

**Figure 1.**
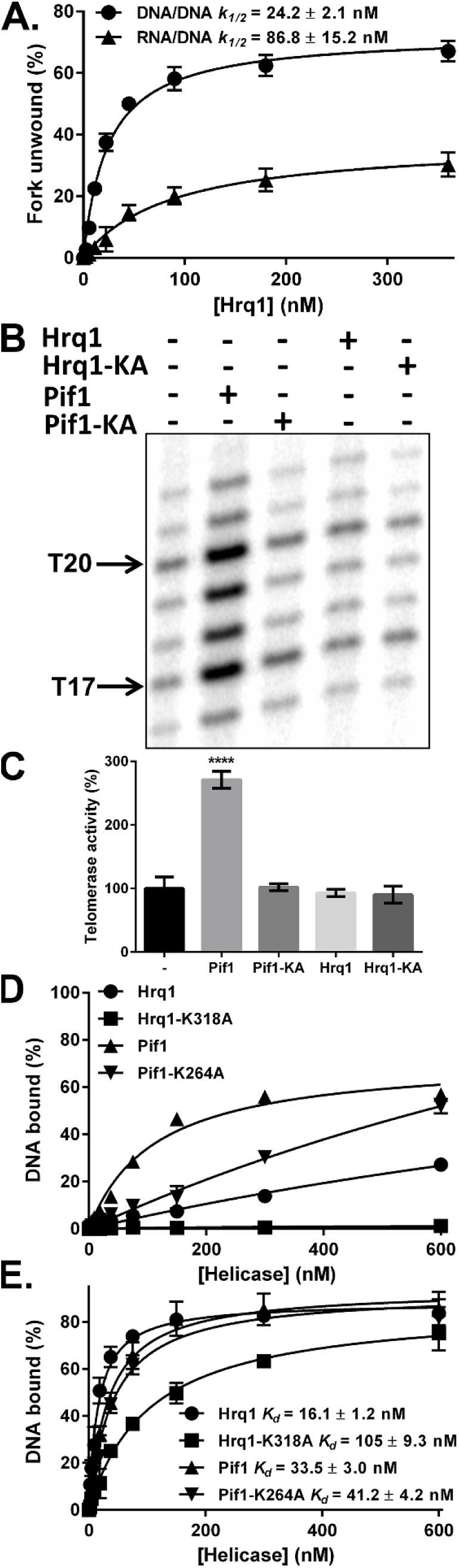
Hrq1 affects telomerase activity by a different mechanism than Pif1. (A) Hrq1 robustly unwinds a fork substrate (25-nt single-stranded DNA (ssDNA) tails, 20-bp DNA-DNA duplex) *in vitro*, but its activity is decreased by an equivalent substrate containing a 20-bp RNA-DNA hybrid duplex. The *k_1/2_* (concentration of Hrq1 needed to unwind 50% of the substrate) values for these curves are listed on the graph. (B) Representative image of *in vitro* telomerase extension of the Tel15 substrate in the absence (−) or presence of 50 nM of the indicated recombinant helicase preparation. Prominent bands are noted at the T17 and T20 positions, indicating that telomerase often stalls after extending Tel15 by +2 and +5 nt. Hrq1-KA and Pif1-KA denote the Hrq1-K318A and Pif1-K264A proteins, respectively. (C) Quantification of overall telomerase activity from the triplicate experiments represented in (B). Telomerase activity was normalized to 100% for the reactions lacking added recombinant helicase. Pif1 significantly increases overall telomerase activity (**** *p* < 0.0001), but the other proteins have no effect, relative to reactions lacking added helicase. (D) Hrq1 and Hrq1-K318A bind poorly to the Tel15 substrate. The plotted data represent the results of gel shift experiments using radiolabeled Tel15 and the indicated concentrations of Hrq1, Hrq1-K318A, Pif1, and Pif1-K264A protein. (E) All tested helicases display better binding to the Tel30 substrate. Gel shifts were performed as in (D) but with a 30-nt substrate. The *K_d_* (concentration of protein needed to bind 50% of the substrate) values for these curves are listed on the graph. In panels A and C-E, all values are the means of ≥3 experiments, and the error bars correspond to the standard deviation (SD). Significant differences were determined by one-way analysis of variance (ANOVA) followed by Tukey's multiple comparison test (*n=* 3).

To begin to determine how Hrq1 affects telomerase activity, we performed a standard *in vitro* telomerase primer extension assay (21) that was previously used to demonstrate the effects of Pif1 on telomerase activity (11). These assays enable one to directly determine the effects of a purified helicase on telomerase activity on a 15-nt telomeric repeat sequence oligonucleotide (Tel15, Table 1), including total activity, extension processivity, and the final distribution of products at each extension position (21,23). Here, we used 50 nM recombinant Pif1 and the catalytically inactive Pif1-K264A mutant as positive and negative controls, respectively, and for comparison with 50 nM recombinant Hrq1 and Hrq1-K318A. Figure 1B shows the seven typical extension products (+1 to +7 or T16 to T22) synthesized by telomerase, and Figure 1C shows the effects of the four recombinant helicase preparations. As previously reported (11), Pif1 increased overall telomerase activity (Fig. 1B,C; Fig. S1A) and altered the final distribution of products and processivity (data not shown), while Pif1-K264A had no significant effects on overall telomerase activity relative to the telomerase alone control (Fig. 1B,C; Fig. S1B). In contrast, neither Hrq1 nor Hrq1-K318A had any effect on overall telomerase activity in this assay (Fig. 1B,C) or when used at concentrations up to 360 nM (Fig. S1C,D). Together these data suggest that Hrq1 and Hrq1-K318A have no effect on telomerase extension of the Tel15 primer.

**Table 1.**
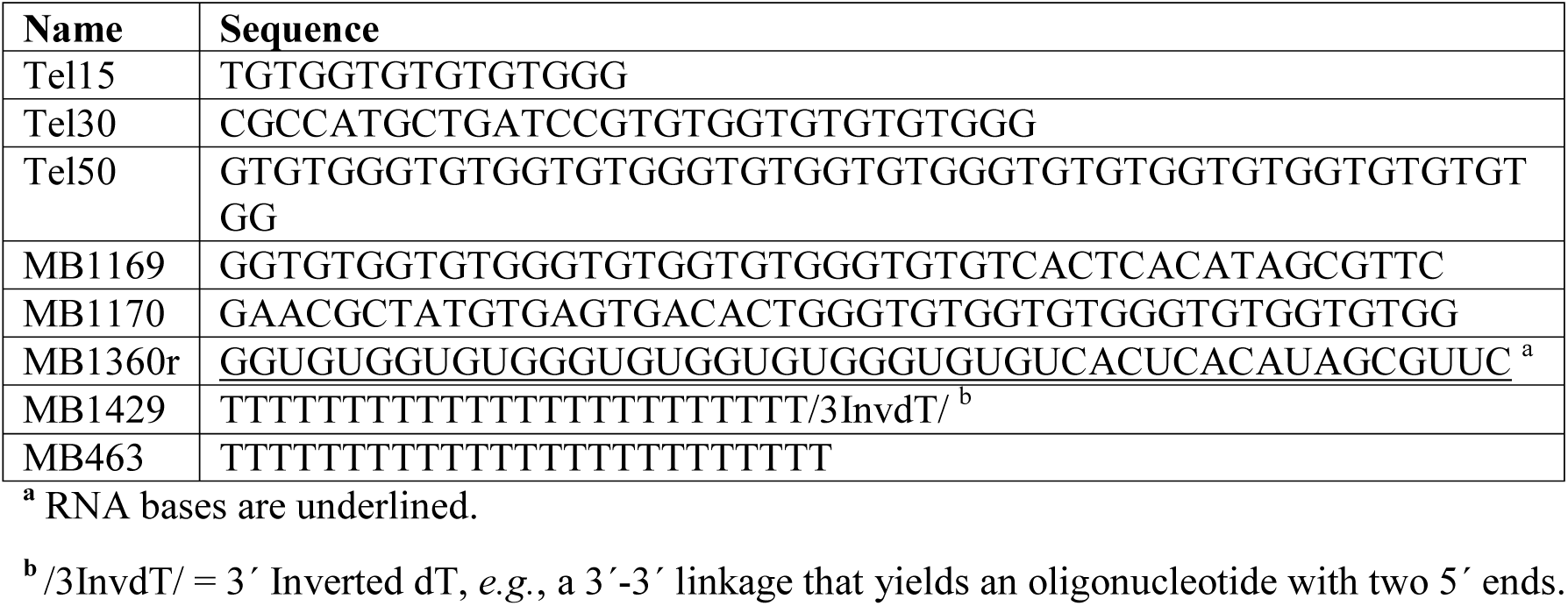
Oligonucleotides used in this study.

Although telomeric repeat sequence ssDNA is a preferred substrate for Hrq1 *in vitro* (15), Hrq1 did not grossly affect telomerase activity in the assays above. We previously reported that Hrq1 requires longer poly(dT) ssDNA substrates for efficient binding (~25 nt (15)), so we reasoned that Hrq1 may require telomeric ssDNA >15 nt to efficiently modify telomerase activity in this *in vitro* system. To test this, we used gel shift assays to observe helicase binding to radiolabeled Tel15 ssDNA. Pif1 and Pif1-K264A both bound to the Tel15 substrate, but Hrq1 binding to Tel15 was much weaker (Fig. 1D). Further, Hrq1-K318A displayed only a basal level of binding to Tel15 (Fig. 1D). The addition of 4 mM ATP to the binding reactions yielded very weak but detectable binding of Hrq1-K318A to Tel15, but this had no significant effect on Hrq1 binding to the primer (data not shown).

Following our finding that Tel15 was too short for efficient binding by Hrq1 and Hrq1-K318A, we next tested binding to a longer 30-nt Tel30 substrate (Table 1). As with Tel15, Pif1 and Pif1-K264A also bound well to Tel30, with *K_d_* = 33.5 and 41.2 nM, respectively (Fig. 1E). Similarly, Hrq1 and Hrq1-K318A both bound more tightly to Tel30 than Tel15, with *K_d_* = 16.1 nM and 105 nM, respectively (Fig. 1E). These results demonstrated that Hrq1 and Hrq1-K318A bound the Tel30 substrate with similar affinities to Pif1 and Pif1-K264A, suggesting that the effects of Hrq1 on *in vitro* telomerase activity should be analyzed using primers longer than the 15-nt Tel15.

### Hrq1 has only subtle effects on telomerase activity on a 30-nt substrate

Based on the above gel shift results, we repeated the telomerase assays with the Tel30 primer. This primer has a random 15-nt 5□ sequence followed by a 15-nt 3□ telomere repeat sequence identical to Tel15 (Table 1). Thus, these primers each feature a 3□ TGGG sequence to pair the primer and *TLC1* RNA template at a single reading frame, allowing us to measure telomerase activity parameters as described for Tel15 (21). By titrating Pif1 into the Tel30 telomerase extension reaction, we found a 1.8-fold increase in total telomerase activity (Fig. 2A), significant changes in the distribution of products at the T31 and T35-T37 extensions (*p*<0.01, Fig. S2A), and a significant decrease in processivity at all positions at high Pif1 concentration (*p*<0.01, Fig. S2B), consistent with previous reports (11) and our results with Tel15 (Fig. 1B,C; Fig. S1A; and data not shown). As observed with Tel15, Pif1-K264A caused no significant effect on total telomerase activity on Tel30 (Fig. 2B), though a slight increase in the distribution of products at T34 (no helicase *vs*. 250 nM, *p*<0.01, Fig. S2C) and subtle alterations in processivity at extensions T32, T34, and T35 were detected at high Pif1-K264A concentrations (*p*<0.01, Fig. S2D).

**Figure 2.**
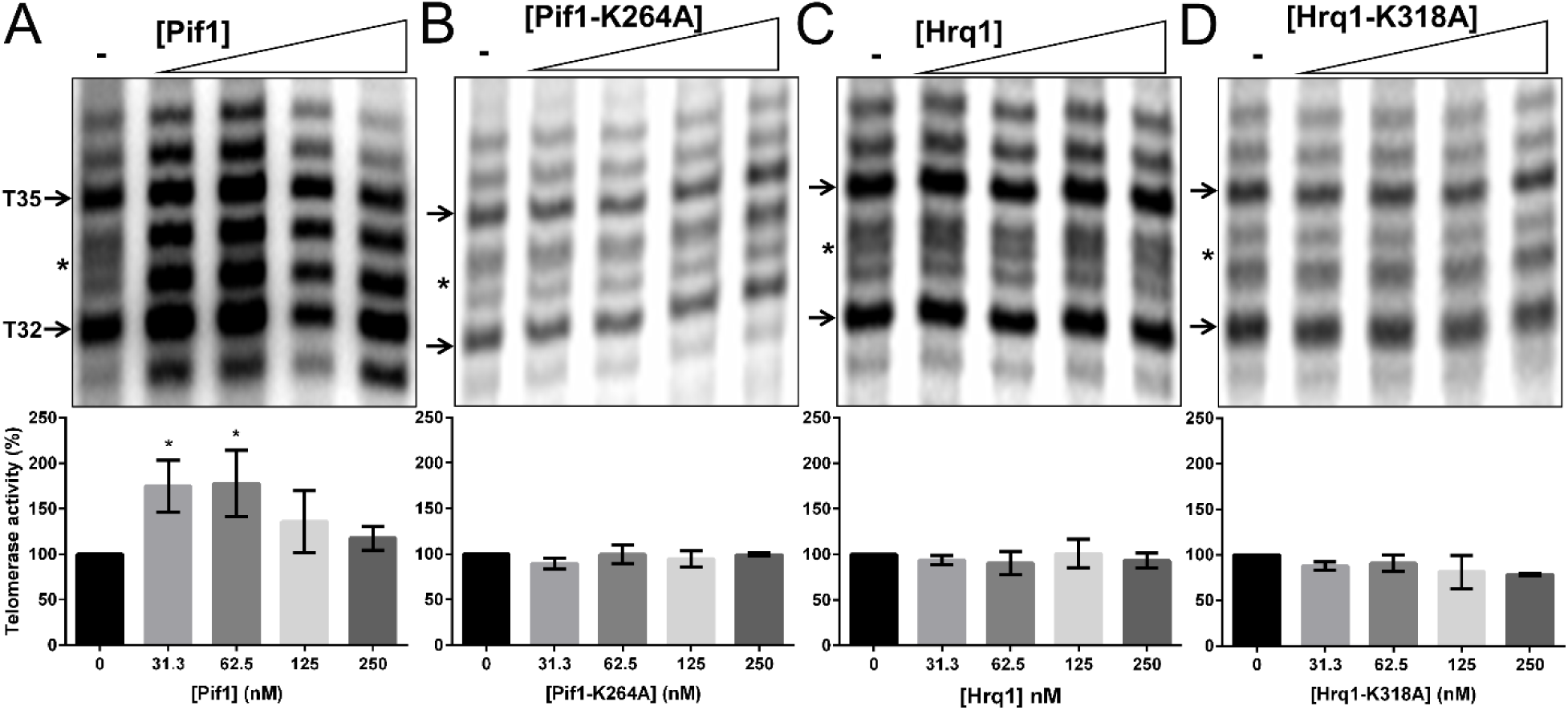
Hrq1 does not affect total telomerase activity on a 30-nt substrate. *In vitro* telomerase primer extension reactions were performed with the Tel30 substrate in the absence of added helicase (−) or presence of increasing concentrations of recombinant Pif1 (A), Pif1-K264A (B), Hrq1 (C), or Hrq1-K318A (D). The upper panels show a representative gel image of experiments performed in triplicate. As with the Tel15 substrate, prominent +2 (T32) and +5 (T35) bands are noted with arrows. The asterisks to the left of each gel image denote putative telomerase stuttering (see Discussion). The lower panels show the quantification of overall telomerase activity from the triplicate experiments. Telomerase activity was normalized to 100% for the reactions lacking added recombinant helicase. The values depicted are the means, and the error bars represent the SD. Pif1 significantly increases telomerase activity by ~1.8-fold when added at 31.3 and 62.5 nM (* *p*<0.01). Significant differences from reactions lacking added helicase were determined by one-way ANOVA followed by Tukey’s multiple comparison test (*n*=3).

Next, Tel30 telomerase assays were performed with increasing concentrations of Hrq1 or Hrq1-K318A added to the reactions. Once again however, no significant effects on total telomerase activity were detected (Fig. 2C,D). Similarly, only minor alterations in processivity and the distribution of Tel30 extension products were found (Fig. S2E-H). These results were unexpected considering the improved binding of Hrq1 and Hrq1-K318A to the Tel30 substrate relative to Tel15 (Fig. 1D,E), and we considered two possible explanations. First, the standard *in vitro* telomerase assay may simply not be sensitive enough to detect the effects of Hrq1 on telomerase. Indeed, the initial *in vivo* observation of Hrq1’s effect on telomere maintenance came from the GCR assay (14), which can report rare events that occur at ~1 × 10^−10^ /generation in *S. cerevisiae* (24). Second, telomerase docks onto the Tel15 and Tel30 substrates at the terminal 3□ TGGG sequence, which may pose a problem for observing effects by a 3□-5□ helicase like Hrq1. The structure of Hrq1 is unknown, though electron microscopy suggests that active recombinant Hrq1 may form a toroid (14,15). Thus, Hrq1 may have to thread onto ssDNA from a free 3□ end. If telomerase is already bound to the 3□ TGGG, this may prevent Hrq1 binding and prohibit Hrq1 from affecting telomerase activity. These possibilities are not mutually exclusive, and both are investigated below.

### Hrq1 increases telomerase activity in a displacement assay

We first addressed the sensitivity of the *in vitro* telomerase primer extension assay using a telomerase displacement assay. In these reactions, telomerase is first allowed to bind to and extend one primer, and then a second primer (bait) of a different size is added after a short incubation time (11,25). This assay has been used for the detection of telomerase displacement from one primer to another by Pif1, which can be monitored by effects on telomerase processivity from each primer in comparison to no helicase controls. Here, we performed two sets of telomerase displacement experiments with either Tel15 or Tel30 first incubated with helicase and/or telomerase for 10 min prior to addition of equimolar bait primer (*i.e*., Tel15 first-Tel30 bait and Tel30 first-Tel15 bait). Results of these experiments are shown in Figures 3 and S3.

**Figure 3.**
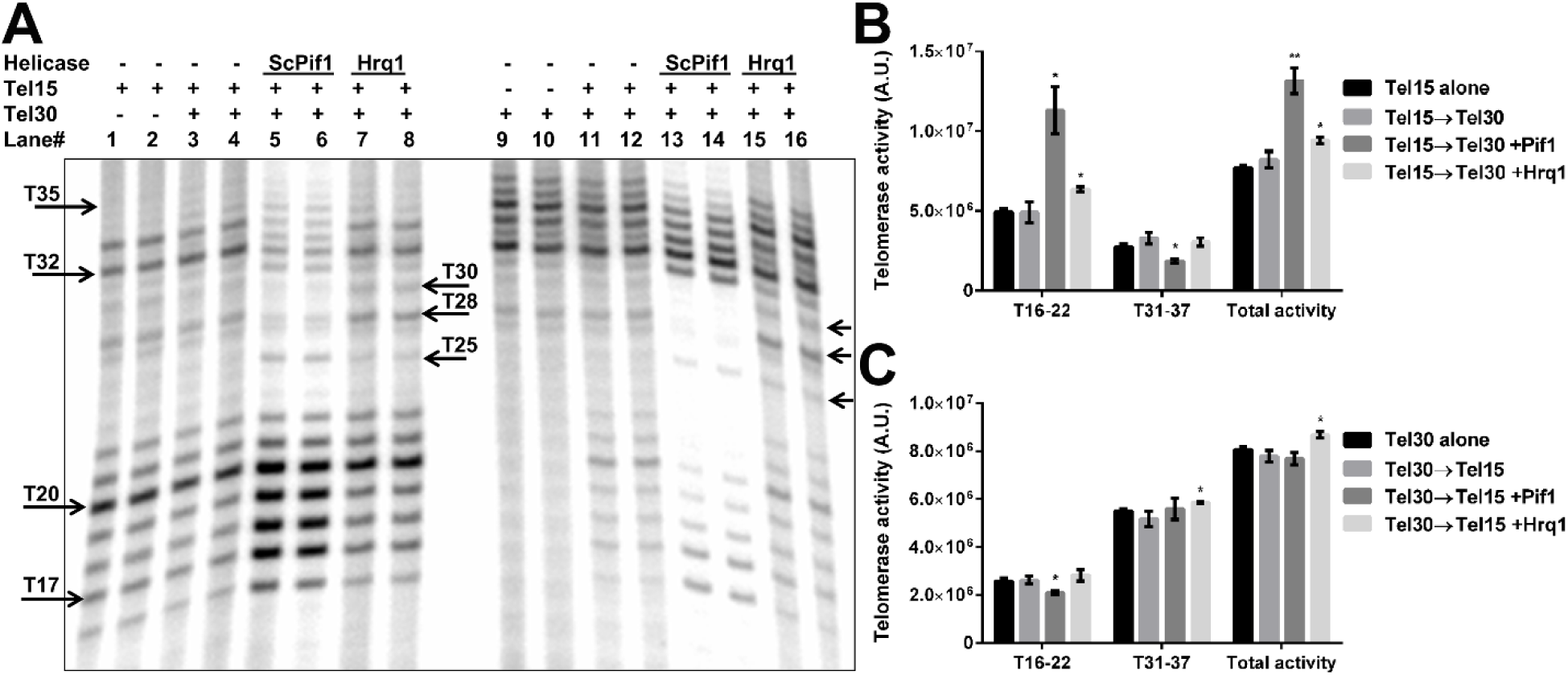
Hrq1 increases telomerase activity in a primer displacement assay. (A) Primer displacement assays that involve incubation of telomerase and 360 nM recombinant helicase with either Tel15 (left, lanes 5-8) or Tel30 (right, lanes 13-16) first, followed by the addition of a second bait substrate (Tel30, left; Tel15, right). Control reactions show the activity of telomerase on Tel15 (lanes 1 and 2) or Tel 30 (lanes 9 and 10) alone, as well as on Tel15 followed by the addition of Tel30 (lanes 3 and 4) or on Tel30 followed by Tel15 addition (lanes 11 and 12) in the absence of added helicase. The generally prominent +2 and +5 extension positions are marked with right-facing arrows for Tel15 (T17 and T20) and Tel30 (T32 and T35). The left-facing arrows denote extension positions T25, T28, and T30, labeled in the middle of the image. Duplicate reactions are shown in neighboring lanes for each set of conditions. (B) Quantification of telomerase activity in arbitrary units (A.U.) on Tel15 extensions (T16-T22), Tel30 extensions (T31-T37), and total activity for reactions performed as shown in lanes 1-8 in (A). Tel15 alone denotes reactions such as those in lanes 1 and 2, Tel15→Tel30 denotes reactions such as those shown in lanes 3 and 4, Tel15→Tel30 +Pif1 denotes reactions such as those shown in lanes 5 and 6, and Tel15→Tel30 +Hrq1 denotes reactions such as those shown in lanes 7 and 8. * *p*<0.01 and ** *p<* 0.001 *vs*. the Tel15→Tel30 reactions. (C) Quantification of telomerase activity as in (B) for reactions performed as shown in lanes 9-16 in (A). Tel30 alone denotes reactions such as those in lanes 9 and 10, Tel30→Tel15 denotes reactions such as those shown in lanes 11 and 12, Tel30→Tel15 +Pif1 denotes reactions such as those shown in lanes 13 and 14, and Tel30→Tel15 +Hrq1 denotes reactions such as those shown in lanes 15 and 16. * *p*<0.01 vs. the Tel30→Tel15 reactions. In (B) and (C), the data are the means of three independent experiments, and the error bars represent the SD. Significant differences from reactions lacking added helicase were determined by multiple *t* tests using the Holm-Sidak method, with α = 5% and without assuming a consistent SD (n=3). There were no significant differences in reactions containing one DNA substrate *vs*. two in the absence of added helicase.

In Figure 3A, lanes 1-4 and 9-12 are no helicase controls with (lanes 3, 4, 11, and 12) or without (lanes 1, 2, 9, and 10) the bait primer added. When telomerase was incubated with Tel15 first, the addition of Tel30 as bait had no effect on overall telomerase activity on Tel15, and only a slight amount of extension from Tel30 was observed (Fig. 3A,B, lanes 1-4). Subtle changes were also detected in the processivity of telomerase on the Tel15 substrate (*p*<0.01, Fig S3B). This confirmed that the majority of telomerase bound to the first primer with little evidence of dissociation and binding to the bait primer. Telomerase is able to synthesize long DNA additions up to hundreds of nucleotides by a set of dual processes that are sometimes referred to as nucleotide addition (Type I) and repeat addition processivity (Type II) (26). The two types of addition are a consequence of short RNA telomerase template regions ranging from 8-31 nt. Type I processivity refers to the ability of telomerase to add nucleotides up to a limit enforced by the finger subdomain of telomerase (27,28). To extend DNA beyond this limit, telomerase must either dissociate from the template or reposition the active site on the template for further elongation (Type II). Failure to dissociate from the initial primer suggests that telomerase is limited to Type I processivity in the absence of helicase activity.

The addition of Pif1 to the dual primer assay significantly reduced Type II processivity while increasing extension from Tel15 (2.3-fold) (Fig. 3A,B). Increased Type I extension activity from Tel30 is partially masked during quantification by the decrease in the background signal we observe in telomerase reactions containing Pif1 (Fig. 3A, compare lanes 3-4 with lanes 5-6). As previously reported, Pif1 also significantly decreased processivity and altered the signal distribution of extensions from both primers, limiting most extensions to the first 5 nt of each primer (Fig. S3A,B). Pif1 also caused a strong stop during the first round of Type II processivity (extensions T18-T29) at T25 but inhibited stops at T28 and T30 (Fig. 3A). These results confirm the effectiveness of this assay for measuring telomerase displacement.

When repeating the Tel15-to-Tel30 primer displacement assay with Hrq1, we found a significant increase in total telomerase activity (1.3-fold, *p*<0.01) (Fig. 3A lanes 7-8, Fig. 3B) that was specifically due to increased activity on Tel15 (Fig. 3B). Because telomerase can efficiently extend Tel15 to position T34 in the absence of Tel30 or helicase (Fig. 3A lanes 1-2), some of the Tel30 extension signal comes from telomerase Type II extension. While this complicates measurements of processivity (Fig. S3B), this observation suggests that Hrq1 cannot efficiently displace telomerase from short primers like Tel15, consistent with our results shown in Figures 1B,C and S1C. Hrq1 also caused a stalling event at T25, but this effect was weaker than that observed with Pif1 (Fig. 3A lanes 5-8). Unlike Pif1, however, Hrq1 caused an increase in the amount of signal (*i.e*., decreased processivity) at positions T28 and T30 (Fig. 3A). Again, there were no significant effects on the signal distribution of extension products (Fig. S3A), but decreases in processivity were found at extension products T33 (*p*<0.001), T35, and T36 (both *p*<0.01, Fig. S3B). Together, these data suggest that Hrq1 is weakly able to displace telomerase from Tel30, enabling subsequent productive binding of telomerase to Tel15. Hrq1 caused strong stops after primer extension reached ~24 nt, suggesting some interaction with telomerase on longer ssDNA substrates.

Unlike dual primer experiments with Tel 15 added first and Tel30 used as bait, when Tel30 was added to reactions before Tel15 as the bait, a small amount of telomerase activity on the bait primer was evident in the absence of added helicase (Fig. 3A, compare lanes 9-10 with 11-12). Under these conditions, the addition of Pif1 had no significant effect on total telomerase extension activity (Fig. 3C), though there was a small reduction in telomerase extension activity from Tel15 (*p*<0.01) (Fig. 3A lanes 13-14, Fig. 3C). As observed in the previous assays, the clearing of background by Pif1 combined with the reduced read-through from telomerase made it difficult to parse increased Type I from decreased Type II processivity effects (Fig. 3A, lanes 11-14). The extension products from each substrate were primarily limited to the first 5 nt (Fig. 3A, lanes 13-14, Fig. 3C, S3C), similar to observations with Tel30 as bait. This is reflected in the fact that processivity was significantly reduced at all but one extension product from Tel15 (*p*<0.01-0.0001) and all extensions from Tel30 (*p<*0.0001) (Fig. S3D). A strong stop at T25 was also observed with Pif1 that was not seen in the no helicase controls (Fig. 3A lanes 13-14). These data support previous reports that Pif1 actively displaces telomerase from Tel15 or Tel30, primarily limiting extension to the first 4-5 nt beyond the primer ends (11). Pif1 appears to inhibit both Type I and Type II telomerase processivity.

Hrq1 caused a small but significant (p<0.01) increase in total telomerase activity when Tel30 was added before the Tel15 bait (Fig. 3A lanes 15-16, Fig. 5C). Alterations in the signal distribution at T17, T21, T22, and T35 (Fig. S3C), as well as in processivity at T17, T19, T20, T33, and T34 (Fig. S3D), were found. Strong stops were observed at positions T25, T28, and T30, suggesting that the presence of Hrq1 is increasing telomerase stalling during the second set of extensions on Tel15 (T23-T30). Together these results further indicate that Hrq1 only effects telomerase activity on longer (>15 nt) substrates. They also suggest that the standard telomerase primer extension assay is not optimal for detecting the effects of Hrq1 on telomerase activity.

### Hrq1 and Pif1 display opposite effects on telomerase extension of a long (50-nt) primer

Having investigated the sensitivity of our telomerase primer extension assay, we next sought to determine if Hrq1 requires a free 3□ ssDNA for loading and binding by performing gel shift assays with recombinant Hrq1 and a standard 25-nt poly(dT) substrate or one containing an inverted 3 terminal base. This modification is introduced by a 3□-3□ linkage, yielding an oligonucleotide substrate with two 5 ends and, thus, no free 3 end. As shown in Figure S4A, Hrq1 displayed nearly indistinguishable binding (p=0.94) to both 25-nt substrates, with binding constants of ~35 nM. Therefore, Hrq1 can internally bind to ssDNA without the need for a free 3 end, which also corresponds to the ability of Hrq1 to bind ssDNA bubble substrates *in vitro* (15).

Regardless, the subtle *in vitro* effects of Hrq1 on telomerase activity we presented above contrast with genetic and biochemical evidence of its potential role in telomere homeostasis (14,15). Therefore, primer design remained a concern. The Tel15 and Tel30 primers were designed to accurately align the *TLC1* template RNA with telomeric repeat ssDNA using a TGGG sequence at the 3 end of the oligonucleotides (Table 1). This feature enables primer extension from a single frame, allowing label correction for measurement of telomerase processivity (11,29). As a consequence, though, telomerase loading at the 3□ end of the ssDNA could preclude Hrq1 from functional interactions with an active telomerase-ssDNA complex if Hrq1 binds the 5□-end or central portion of the substrate. Data from our telomerase assays and binding assays do not support a mechanism where Hrq1 exclusively removes telomerase bound at telomere 3□ ends, indicating that Hrq1 can bind upstream from telomerase. Such a situation would establish a futile catalytic cycle where Hrq1 translocates 3□-5□ away from telomerase without ever affecting its activity. For Hrq1 to have a measurable effect on telomerase activity *in vitro*, telomerase may require an internal binding site on a long ssDNA substrate. In this scenario, telomerase complexes loaded or stalled at sites distal to the 3□ ssDNA end could be removed or disrupted by the 3□-5□ helicase activity of Hrq1.

To test this hypothesis, we repeated telomerase assays with a Tel50 substrate that was all telomeric repeat sequence (*i.e*., lacking 5□ random sequence as in Tel30) and without a 3□-TGGG guide sequence, thus encouraging telomerase binding to a more central portion of the substrate. Without the guide sequence, we cannot determine telomerase processivity or the distribution of radiolabeled extension productions, but total telomerase activity can still be measured. Other caveats with longer telomeric primers included a reduced yield of extension products and increased nuclease activity by our telomerase-enriched extract. Nuclease activity is known to be associated with telomerase from all examined sources, including *S. cerevisiae* and *H. sapiens* (21,30-33). This activity is specific, proportional to telomerase activity, and can be reconstituted in telomerase/*TLC1*-generating rabbit reticulocyte lysates (34,35).

In initial experiments using Tel15, Tel30, or Tel50 primers, we noted a characteristic pattern of RNaseA-sensitive products that were shorter than the Tel30 and Tel50 substrates (Fig. S4B). In the case of Tel50, shorter products appeared to be initiated from a cleavage event between nucleotides 21 to 25 of the Tel50 primer, followed by telomerase extension (Fig. S4C and data not shown). Extension products were seen at T26, the first labeled position following nuclease cleavage, independent of the length of primer used. Blocking the 3□ end of Tel50 with terminal transferase and ddATP (creating the Tel50B substrate) inhibited direct telomerase extension from Tel50 (p<0.0001), but nuclease-cleaved extension products were not inhibited (Fig. S4C,D). We used this phenomenon as a tool to examine the effects of helicase titrations on telomerase activity from primers > 30 nt. Direct Tel50 extension products and the nuclease-cleaved extension products were quantified. Direct Type I extensions from the 3□ end of Tel50 were not well resolved, so they were quantified as one unit (T51-58). Due to this, all processivity and signal distribution data were only calculated based on nuclease-cleaved extension products (approximately T26-32).

Results of assays containing Pif1 showed a complex activity profile over the range of concentrations tested (Fig. 4A,B and S5A,B). In the absence of Pif1, 73% of the signal was in the nuclease-cleaved extension products (T26-32), while 27% was in the direct extension products (T51-58). Telomerase extension from the cleaved Tel50 substrate yielded seven bands, typical for most reported Type I activity (Fig. 4A). The distribution of radiolabel, however, was atypical relative to Tel15 and Tel30 assays because the +4 and +7 extension products, rather than +2 and +5, displayed the highest levels of signal (Fig. 4A). This observation could be due to extension from either the 5□ or 3□ nuclease cleavage fragments, which can both act as substrates (31). Accumulation of signal at position T32 suggests that telomerase stalls at this positon. Reduced Type II processivity is reflected in the minimal amount of extension observed between T33 and T50. At lower concentrations (up to 90 nM), Pif1 significantly stimulated total telomerase activity (*p*<0.01), but telomerase activity levels decreased to below the no helicase control at higher concentrations of added helicase (*p*<0.01; Fig. 4A,B). Indeed, at 360 nM Pif1, total telomerase activity was significantly decreased compared to the no helicase control (*p*<0.001, Fig. 4A,B). To our knowledge, this biphasic concentration-dependent switch from activator to inhibitor of total telomerase activity has not been reported, though the inhibition of telomerase activity by Pif1 appeared to directly correlate with the amount of cleaved and extended products as previously observed (31).

**Figure 4.**
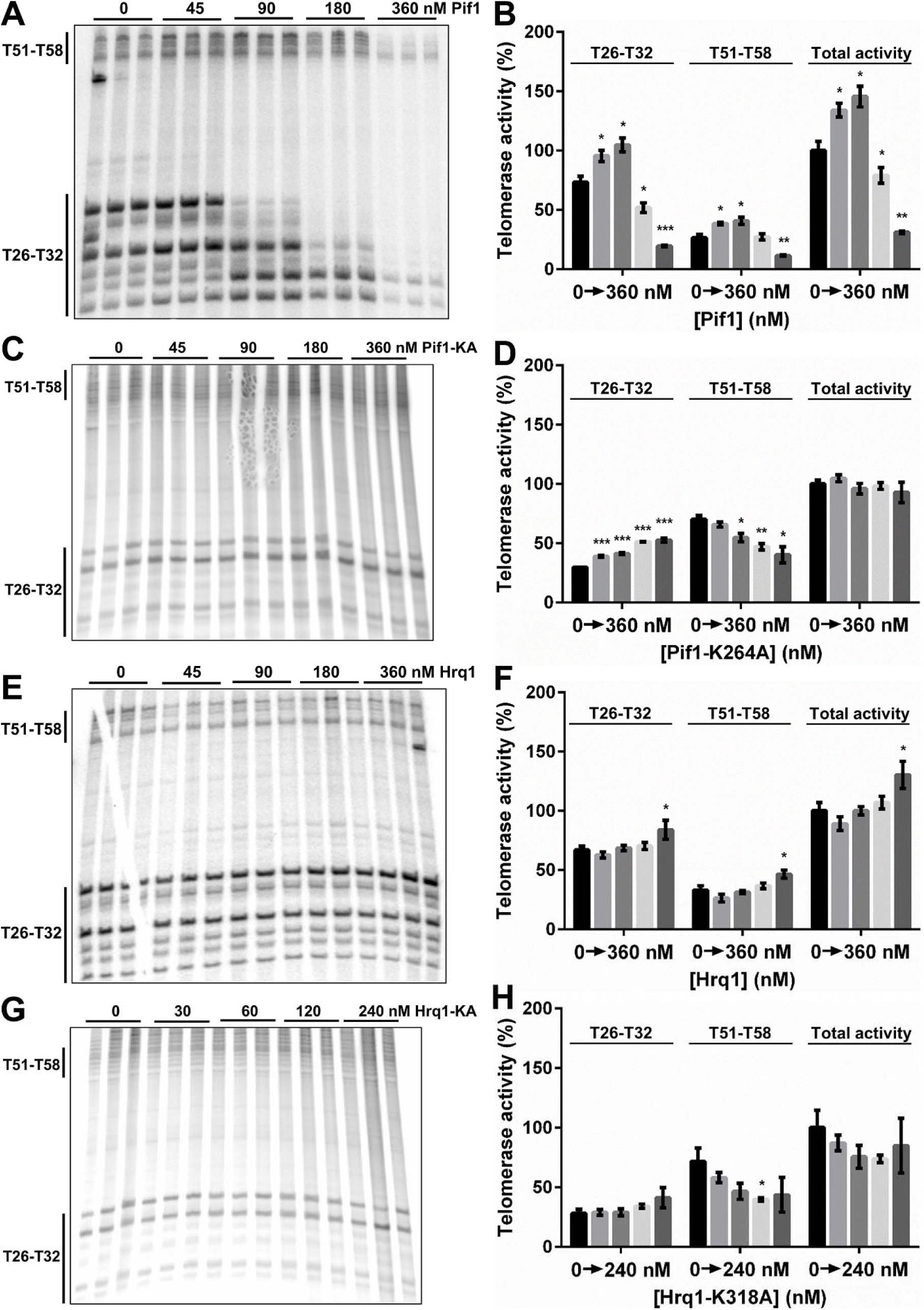
Pif1 and Hrq1 display opposite effects on telomerase extension of the Tel50 substrate. Representative gel images (left) and quantification (right) of telomerase activity on the Tel50 substrate in the absence of added helicase and presence of the indicated concentrations of Pif1 (A,B), Pif1-K264A (C,D), Hrq1 (E,F), or Hrq1-K318A (G,H). The T26-T32 bands are extensions of nuclease-cleaved Tel50, and T51-T58 are direct extensions from the 3ʹ end of Tel50. Total activity refers to the total amount of signal in the lane (T26-T58). The graphed data are the means of three independent experiments, and the error bars represent the SD. Significant differences were determined by multiple *t* tests using the Holm-Sidak method, with α = 5% and without assuming a consistent SD (*n*=3). * *p*<0.01, ** *p<*0.001, and *** *p*<0.0001; all comparisons were made against the reactions lacking added helicase.

Similar assays were next performed with Pif1-K264A (Fig. 4C,D and S5C,D) to determine the effect of Pif1 catalytic activity on telomerase activity with the Tel50 substrate. There was a decreasing trend in direct extension activity from Tel50, but extension from nuclease cleavage products significantly increased (p<0.0001) at all concentrations of Pif1-K264A tested (Fig. 4C). These opposing effects yielded no significant change in total telomerase activity (Fig. 4C), and the radiolabel distribution and processivity effects observed with Pif1 (Fig. S5A,B) were largely absent with Pif1-K264A (Fig. S5C,D). Together, these results indicate that the biphasic stimulation-then-inhibition of telomerase activity by Pif1 (Fig. 4A,B) requires catalytic activity by the helicase, but Pif1-K264A still exerted a non-catalytic effect on both direct extension of Tel50 and extension of nuclease-cleaved Tel50 by telomerase (Fig. 4C,D).

Telomerase assays with Tel50 and increasing concentrations of Hrq1 (Fig. 4E,F and S5E,F) revealed an opposite trend relative to Pif1. Lower concentrations of Hrq1 slightly inhibited telomerase activity (*p*=0.07), but significant stimulation occurred (up to 1.3-fold, *p*<0.01) at higher Hrq1 concentrations (Fig. 4E,F). Similar to Pif1, however, most of the radiolabel signal was found in the extension products at T26-T32 rather than T51-T58 (Fig. 4F). Signal was also noted in the region of Type II telomerase extensions T33-T50 (Fig. 4E). As we observed with Pif1, the ratio of direct to nuclease-cleaved extension products was relatively constant over the range of helicase concentrations tested. Processivity effects were again largely absent, except for decreases at positions T28 and T29 (Fig. S5F). The distribution of radiolabel signal reflects this effect, with more signal in extensions T31 and T32 (Fig. S5E). Overall, as we observed with the Tel15 and Tel30 substrates, Hrq1 acted as a weak but significant activator of telomerase in assays with Tel50.

We also performed Tel50 extension assays with increasing concentrations of Hrq1-K318A (Fig. 4G,H and S5G,H). In this case, however, activity was largely unaltered at the concentrations of helicase tested (Fig. 4G,H). Similarly, only minimal changes in telomerase processivity and signal distribution were observed with Hrq1-K318A (Fig. S5G,H). Therefore, the significant stimulation of telomerase activity by Hrq1 noted above required catalytic activity by the helicase.

### Hrq1 and Pif1 can bind to the same telomeric ssDNA substrate

One curious aspect of the Tel50 telomerase assays above was the biphasic activity curves generated by both Pif1 (Fig. 4B) and Hrq1 (Fig. 4D). In the case of Pif1, total telomerase activity increased to 146% at 90 nM and then decreased in a concentration-dependent manner to 31% at 360 nM Pif1, relative to no helicase controls (Fig. 4B). In contrast, with Hrq1, total telomerase activity decreased to 89% of controls at 45 nM helicase and then increased in a concentration-dependent manner to 130% activity at 360 nM (Fig. 4F). This observation led us to consider the possibility that Pif1 and Hrq1 utilize their equal-but-opposite activities together to maintain telomere length homeostasis *in vivo*. Indeed, chromatin immunoprecipitation demonstrates that Hrq1 (14) and Pif1 (36) can each bind to telomeres *in vivo*. Thus, in cells, Hrq1 may directly interact with Pif1 to alter telomerase activity or possibly even displace Pif1 (or other telomere binding proteins) from telomeric DNA.

Thus, we sought to determine if these helicases act synergistically to affect telomerase activity on the same substrate *in vitro* by two approaches. First, to investigate if dual helicase binding was possible, we used radiolabeled Tel50 substrate and performed agarose gel shift assays with recombinant Hrq1 and/or Pif1. We observed a super-shift of Tel50 when the concentration of Pif1 was held constant and increasing concentrations of Hrq1 were added and *vice versa* (Fig. S6A-D). These results could be due to either both Hrq1 and Pif1 binding to the same Tel50 substrate or to direct protein-protein interactions between Hrq1 and Pif1, one of which is substrate bound. Although we cannot exclude the latter, it should be noted that based on the known binding site sizes for Hrq1 (25-30 nt; (15)) and Pif1 (~5 nt; (13,37)), Tel50 is of sufficient length to support binding of both helicases. Therefore, we next performed Tel50 telomerase extension assays in the presence of both Hrq1 and Pif1.

### Combinations of wild type and inactive Hrq1 and Pif1 alternately inhibit and stimulate in vitro telomerase activity

As we observed in Tel50 extension assays with Pif1 alone, telomerase activity first increased and then decreased in a biphasic manner centered at 45 nM when equimolar concentrations of Pif1 and Hrq1 were added (Fig. 5A,B). Unlike with Pif1 alone, however, there was no spike in total activity at 45 and 90 nM helicase for primary extensions from Tel50 (Fig. 5B). This increase was instead observed at the T26-T30 extensions but not in the T51-T58 or T33-T50 extension regions (Fig. 5A,B). A prominent new band was observed at position T26 with 45 nM of each helicase, which was also subject to inhibition as helicase concentrations were increased (Fig. 5A). Comparison of quantified telomerase activity indicated that there is a significant difference (p<0.01) in the extent of inhibition between Pif1 alone and Pif1 + Hrq1 (Fig. 5C). Despite the 1.3-fold increase in total telomerase activity observed with 360 nM Hrq1 alone, when both helicases are present at 360 nM, total extension activity decreased 6.7-fold compared to 3.2-fold with Pif1 alone (Fig. 5C). The presence of Hrq1 also altered the distribution of signal and telomerase processivity compared to assays with Tel50 and Pif1 alone (Fig. S5A,B and S7A,B). These data strongly support the hypothesis that Hrq1 acts synergistically with Pif1, improving the telomerase inhibition activity of Pif1 alone.

**Figure 5.**
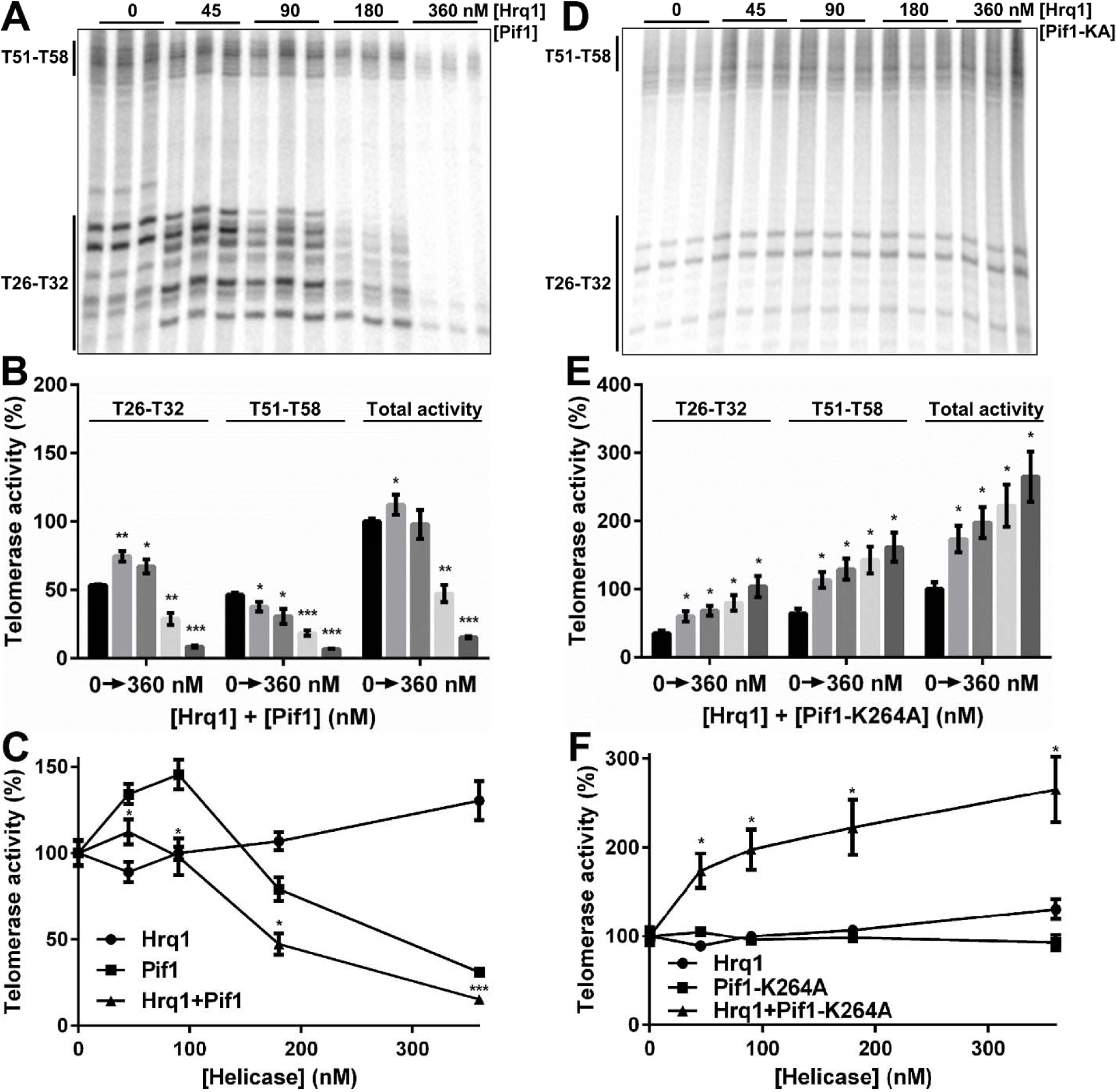
Combinations of wild type and inactive Hrq1 and Pif1 alternately inhibit and stimulate *in vitro* telomerase activity. Representative gel images (top), quantification of telomerase activity (middle), and comparisons of overall telomerase activity on the Tel50 substrate (bottom) in the absence of added helicases and presence of the indicated concentrations of Hrq1 and Pif1 (A,B) or Hrq1 and Pif1-K264A (D,E). Equimolar concentrations of helicase were added to each reaction, and the reported concentration is of each helicase (*e.g*., 45 nM is 45 nM each of Hrq1 and Pif1 for a total helicase concentration of 90 nM). The T26-T32 bands are extensions of nuclease-cleaved Tel50, and T51-T58 are direct extensions from the 3ʹ end of Tel50. Total activity refers to the total amount of signal in the lane (T26-T58). The graphed data are the means of three independent experiments, and the error bars represent the SD. In (B) and (E), statistical comparisons were performed relative to the reactions lacking added helicase. In (C), the Hrq1+Pif1 data were compared to Pif1 alone, and in (F), the Hrq1+Pif1-K264A data were compare to both Hrq1 and Pif1-K264A alone. Significant differences were determined by multiple *t* tests using the Holm-Sidak method, with α = 5% and without assuming a consistent SD (*n*=3). * *p*<0.01, ** *p*<0.001, and *** *p*<0.0001.

A caveat of the results described above is that they do not discriminate between the possibility that Hrq1 and Pif1 bind to the same substrate to perform their functions separately or if the helicases physically interact to exert a concerted effect on telomerase. As the catalytic activity of Pif1 is necessary to impact overall telomerase activity on Tel50 (Fig. 4C,D), but Pif1-K264A retains DNA binding activity (Fig. 1D,E), we next tested the combination of Hrq1 and Pif1-K264A in the Tel50 extension assays. As expected, these assays demonstrated that Pif1-K264A does not support the inhibition of telomerase activity (Fig. 6D,E). However, the activation of telomerase activity displayed by Hrq1 alone (Fig. 4E,F) was significantly increased (p<0.01), for both the direct extensions and nuclease-cleaved extensions, in the presence of all concentrations of Hrq1 + Pif1-K264A tested (Fig. 5D-F). This result was unexpected because Hrq1 had never yielded greater than a 1.3-fold increase in total telomerase activity in any of our assays. Despite the increase in total telomerase activity, the presence of Pif1-K264A with Hrq1 eliminated the signal distribution and processivity effects observed with both wild-type helicases together (Fig. S7A,B) or Pif1 alone (Fig. S5A,B). Again, these results supports the hypothesis that Hrq1 and Pif1 act synergistically, rather than additively, to alter telomere extension activity by telomerase.

**Figure 6.**
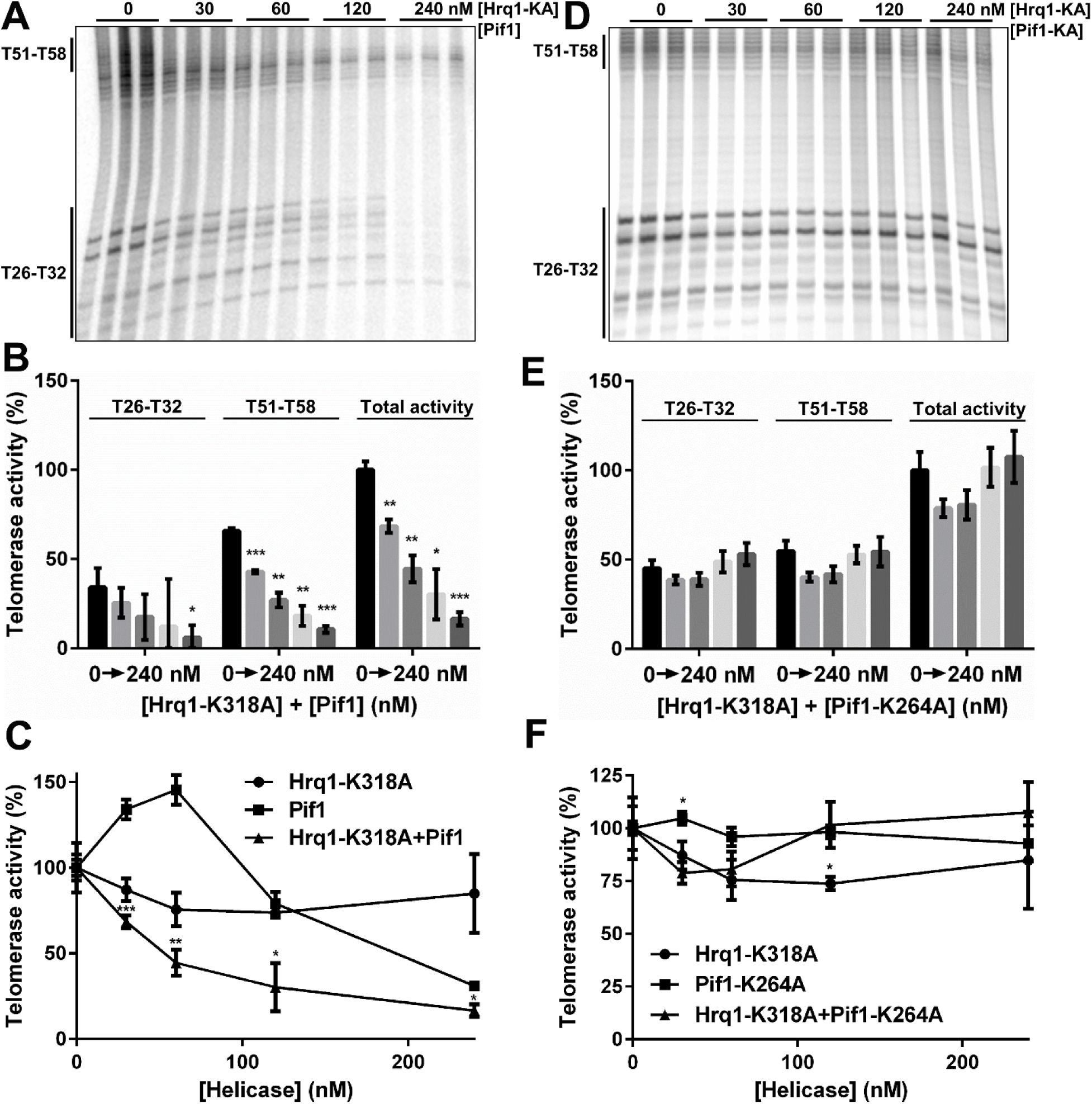
Hrq1-K318A stimulates the inhibition of telomerase activity by Pif1. Representative gel images (top), quantification of telomerase activity (middle), and comparisons of overall telomerase activity on the Tel50 substrate (bottom) in the absence of added helicases and presence of the indicated concentrations of Hrq1-K318A and Pif1 (A,B) or Hrq1-K318A and Pif1-K264A (D,E). Equimolar concentrations of helicase were added to each reaction, and the reported concentration is of each helicase (*e.g*., 30 nM is 30 nM each of Hrq1 and Pif1 for a total helicase concentration of 60 nM). The T26-T32 bands are extensions of nuclease-cleaved Tel50, and T51-T58 are direct extensions from the 3' end of Tel50. Total activity refers to the total amount of signal in the lane (T26-T58). The graphed data are the means of three independent experiments, and the error bars represent the SD. In (B) and (E), statistical comparisons were performed relative to the reactions lacking added helicase. In (C), the Hrq1-K318A+Pif1 data were compared to Pif1 alone, and in (F), the Hrq1-K318A+Pif1-K264A data were compare to both Hrq1-K318A and Pif1-K264A alone. Significant differences were determined by multiple *t* tests using the Holm-Sidak method, with α = 5% and without assuming a consistent SD (*n*=3). * *p*<0.01, ** *p*<0.001, and *** *p*<0.0001.

To perform the reciprocal experiment, we tested the combination of wild-type Pif1 and Hrq1-K318A in the Tel50 telomerase assay. Together, these recombinant proteins acted as a concentration-dependent inhibitor of telomerase activity (Fig. 6A-C). Processivity and the distribution of radiolabel were both altered compared to assays containing Hrq1 + Pif1 (Fig S7E,F *vs*. S7A,B). Hrq1 in combination with Pif1 did not have a drastic effect on processivity compared to Pif1 alone (Fig. S7B *vs*. S5B), but Pif1 + Hrq1-K318A effects on processivity (Fig. S7F) suggest an interaction with Pif1. These results indicate that the catalytic activity of Hrq1 is not necessary to stimulate the telomerase inhibition activity of Pif1, again suggesting a protein-protein interaction between Hrq1 and Pif1.

As a control we also tested telomerase extension of Tel50 in the presence of Hrq1-K318A + Pif1-K264A (Fig. 6D-F). No significant changes in telomerase activity were detected over the range of concentrations tested (Fig. 6E). However, some minor effects on radiolabel distribution and processivity were observed (Fig. S7G,H). Compared to either inactive helicase alone, there was largely no difference in overall telomerase activity when both were combined (Fig. 6F). Thus, the ATPase-null mutants together largely lacked the ability to affect telomerase activity. The subtle effects that were noted could be due to occlusion of the cleavage site(s) and 3ʹ ends of the Tel50 substrate by bound and immobile helicases at relatively high concentration.

## DISCUSSION

### Hrq1 and Pif1 function together to maintain telomere length homeostasis

It has been suggested that Pif1 functions as a telomerase inhibitor by unwinding *TLC1* RNA-telomeric DNA hybrids, thus evicting telomerase from chromosome ends (38). Further, as we (*e.g*., Fig. 2A) and others (11,16,17) have shown, Pif1 requires catalytic activity for this effect. Results from *in vivo* assays also indicate that Hrq1 is a telomerase inhibitor, but *hrq1-K318A* cells do not exhibit the telomere maintenance defects of *hrq1Δ* cells (14). This indicates that catalytic activity by Hrq1 is not necessary for telomerase inhibition, and thus, that its mechanism of inhibition differs from that of Pif1. Indeed, our early results demonstrated that RNA-DNA hybrids inhibit Hrq1 helicase activity (Fig. 1A) rather than stimulate activity as with Pif1 (12,39), so the basic biochemistry of these enzymes also supports mechanistic differences.

Using an *in vitro* telomerase primer extension assay, a similar situation was revealed. At low concentrations, Pif1 stimulated telomerase activity (Fig. 2A and 4A,B), presumably by freeing stalled telomerase complexes for productive rebinding and another round of primer extension (13). However, at increased Pif1 concentrations, total telomerase activity was significantly decreased (Fig. 4A,B). Hrq1 displayed the opposite trend. Low concentrations of the helicase slightly decreased telomerase activity to a repeatable but not statistically significant amount, but total telomerase activity was significantly stimulated in the presence of 360 nM Hrq1 (Fig. 4E,F).

To investigate this inconsistency, we performed telomerase primer extension assays and included both Hrq1 and Pif1 in the reaction together to more closely mimic the situation in cells. Under these conditions, the combined helicases displayed the biphasic telomerase activity stimulation-then inhibition effect observed with Pif1 alone. However, the extent of stimulation was significantly decreased, and the inhibition phase was significantly stronger (Fig. 5A-C). Inclusion of ATPase-null helicases in these assays also addressed the gene deletion *vs*. inactive allele dichotomy observed *in vivo* (14). The combination of Hrq1-K318A and wild-type Pif1 yielded stronger telomerase inhibition than Pif1 alone (Fig. 6A-C), explaining why telomere maintenance in *hrq1-K318A* cells resembles wildtype rather than *hrq1Δ*. Similarly, the combination of Pif1-K264A and wild-type Hrq1 stimulated telomerase activity at all concentrations of helicase tested, which matches the longer telomeres found in *pif1-K264A* cells. This is the first mechanistic insight into the roles of Hrq1 in telomere maintenance, especially in combination with Pif1. At least one of the helicases had to be active to observe these effects though, as reactions containing Hrq1-K318A and Pif1-K264A together did not differ much from reactions containing no helicase or either of the inactive helicases alone. Aside from explaining the apparently contradictory effects of the *hrq1Δ* and *hrq1-K318A* alleles on telomerase (14), these *in vitro* data also highlight the dangers of being too reductive when investigating biological processes by molecular biochemistry. Concentrating on a single player (Hrq1 or Pif1) does not yield the full story, and this is an especially important lesson to remember when dealing with a complex process like telomere maintenance. Future work should include other helicases (*e.g*., Sgs1) and proteins found at telomeres (*e.g*., Cdc13) to truly recapitulate telomere length homeostasis at a mechanistic level.

### Telomeric ssDNA length impacts the activity of Hrq1

Together, our data suggest that Hrq1 and Pif1 interact at telomeres to modulate each other’s biochemical activities, and this synergism affects telomerase activity to maintain telomeres at a normal homeostatic length. Telomere length homeostasis is a key function of all eukaryotic cells, and telomerase is a central enzyme in this process in most cases (40). Balanced against the need for telomerase binding and extension of telomeres is the need to avoid telomere additions at DSBs. Cells must be able to distinguish DSBs from telomeres, and helicases play a critical role in telomere length control and displacement of telomerase from DSBs (14,38). Pif1 and Hrq1 are implicated in both processes, suggesting that the helicases are involved in separate pathways, preparing DNA repair intermediates for homologous recombination (HR), protecting DSBs from telomere additions, and in the maintenance of telomere length.

Pif1 is a central regulator of telomere length and is important for yeast cells to distinguish telomeres from DSBs. Recently, inducible short telomere systems and single telomere extension (STEX) assays have been used to probe the effects on Pif1 binding and activity at telomeres of different lengths (36,41). STEX experiments demonstrate that telomerase processivity and the fraction of telomeres extended *in vivo* increases in the absence of Pif1 (36). Using induced short telomeres of different lengths, the authors showed that preferential binding of Est2 to short telomeres is lost in *pif1-m2* cells. They conclude that Pif1 preferentially binds to long telomeres, allowing short telomeres to be extended. The gel shift data reported here supports this conclusion, with better binding of Pif1 to the Tel30 *vs*. Tel15 substrate (Fig. 1D,E). In addition, the degree of telomerase activity inhibition by Pif1 increased with increasing primer length (Fig. S8).

Using a similar *in vivo* telomere length induction system, another group found a transition point where telomeric sequences of 34-125 bp are recognized as short telomeres and are preferentially extended by telomerase (41). Their model suggests that at chromosome ends of ~40 bp, Pif1 blocks resection and telomerase activity, acting as a checkpoint for distinguishing DSBs from telomeres. The authors argue that at telomeres > 34 bp, Cdc13 recruitment overwhelms Pif1 inhibition, allowing length extension by telomerase.

While our analyses do not directly address the issue of telomere addition at DSBs, gel shifts demonstrate that Hrq1 efficiently binds ssDNA ≥ 25 nt and favors binding to telomeric ssDNA substrates (15) (Fig. 1E). In most stages of the cell cycle, wildtype *S. cerevisiae* telomeres consist of an average of 250-400 bp of TG_1_-_3_ repeats that end in a 10-15 nt 3ʹ G-strand overhang (42). During late S-phase, the ssDNA tail increases in length to approximately 50-100 nt, offering a time-dependent window when Hrq1 could bind directly to ssDNA at telomeres. Pif1, on the other hand, requires only a 5-nt gap to bind to and remove telomerase *in vitro* (13). When one considers that during DSB repair ssDNA can reach tens of thousands of nucleotides in length (43), resected DSB would be an optimal substrate for Hrq1 binding compared with telomeres. This evidence points to Pif1 being the central helicase in telomerase removal from telomeres due to higher likelihood of binding to telomeres. Synergism between Pif1 and Hrq1 to remove telomerase is likely to occur at hyperextended telomeres or, more likely in wildtype cells, at DSBs. Data presented here demonstrate that Pif1 is a more active inhibitor of telomerase activity in the presence of either Hrq1 (Fig. 5A,C) or catalytically inactive Hrq1 (Fig. 6A,C). These results support earlier *in vivo* data indicating that both Pif1 and Hrq1 are essential for avoiding telomere additions at DSBs, while Hrq1 effects on telomerase activity were only observed in the *pif1-m2* background (14).

Multiple protein complexes are involved in binding telomeres and protecting them from nuclease- and helicase-controlled resection, thus preventing activation of DNA damage signaling pathways (41,44). These proteins include MRX (Mre11/Rad50/Xrs2), CST (Cdc13/Stn1/Ten1), Ku70/80, and the Rap1/Rif1/Rif2 dsDNA binding complex (reviewed in (45)). As replication proceeds through G-rich telomeric DNA, these protein complexes must be removed and replaced as the replication fork passes. Failure to coordinate this process can lead to replication fork collapse, arrest of cell growth, or failure to protect the actively dividing chromosome ends. The ability of Hrq1, a weak activator of telomerase, to improve Pif1 inhibition of telomerase indicates that the mechanism of Hrq1 control of telomerase activity is distinct from Pif1. This is confirmed by data presented here showing that Hrq1 helicase activity is inhibited by RNA in the unbound strand (Fig. 1A), whereas RNA:DNA hybrids stimulate Pif1 helicase activity (39). By extension, the synergism displayed by Pif1 and Hrq1 reducing telomerase activity could also play a role in removing other protein complexes at any stage of telomere replication, telomere elongation, C-strand fill in, or during DSB repair.

### Are other helicases also involved in telomere length homeostasis?

One immediate implication of the work presented here is that the human homologs of Hrq1 and Pif1, RECQL4 and hPIF1 (respectively), may function in a similar synergistic manner to modulate telomere length. Indeed, it has been demonstrated that RECQL4 is involved in telomere maintenance (18,19), and likewise, human PIF1 has shown evidence of being a telomerase inhibitor (46). Therefore, future *in vitro* experiments should address this issue using recombinant RECQL4, hPIF1, and human telomerase.

It should also be noted that many other helicases are also known to affect telomerase. In humans, the BLM (47,48) and WRN (49,50) RecQ family helicases are both involved in telomere maintenance (reviewed in (51)). Similarly, the functional homolog of BLM in yeast, Sgs1, is linked to telomere maintenance, though usually in the context of recombination-mediated telomere lengthening in the absence of telomerase (52-54). However, deletion of *SGS1* can lead to *de novo* telomere addition at DNA DSBs if *EXO1* is also deleted (55). This is similar to the increase in telomere addition after GCR events in *hrq1Δ* cells, though under the same conditions only the inactive *sgs1-K706A* allele rather than *sgs1Δ* results in an increase in telomere addition (46% *vs*. 0%; (14)). In that respect, this is the opposite effect seen with *hrq1-K318A* and *hrq1Δ* (4.5% *vs*. 77% telomere additions). Using recombinant Sgs1 and Sgs1-K706A in the *in vitro* telomerase primer extension assay, especially in combination with wild-type and inactive Hrq1 and/or Pif1, will help to shed light on this phenomenon.

### Nuclease activity associated with telomerase and the impact of DNA helicases

Telomere cleavage by a nuclease activity that is tightly associated with telomerase has been previously reported in yeast (31). This nuclease activity is RNaseA sensitive, is dependent upon telomerase extension, and has not been separated away from telomerase using extensive purification methods. In addition, this activity can be reconstituted with Est2/*TLC1* prepared in rabbit reticulocyte lysates. Niu *et al*. (2000) argue that coupled nuclease cleavage followed by telomere extension plays an important role in yeast, allowing stalled or internally bound telomerase to restart extension, offering an alternative mechanism to Pif1 removal of telomerase (31). The telomere cleavage products we noted appeared independent of the length of the initial oligonucleotide used, with the strongest signals at T27, T31, and T32 (Fig. S4B). Our data support ssDNA cleavage by a tightly coupled nuclease followed by telomerase extension of one or both available 3□ ends. Pif1 alone, or Pif1 and Hrq1 together, reduced both cleavage and extensions in almost equal proportions. These observations suggest a model where low levels of helicase allow more cleavage and extension of telomeres, while high helicase levels block both activities, setting a limit on telomere length and establishing a regulatory mechanism to maintain telomere length homeostasis.

Nuclease cleavage by telomerase is reminiscent of a mode of regulated telomere shortening referred to as telomere rapid deletion (TRD) in both human a mouse cells (56). This telomere trimming phenomenon is thought to allow resolution of structured DNA (T-loops, G-quadraplex, and HR intermediates) that form at telomeres due to G-rich 3□ overhangs, culminating in the release of extrachromosomal telomeric DNA. While our data cannot distinguish between these models, increasing helicase activity should reduce both TRD and nuclease cleavage to free stalled telomerase. In the case of TRD, any increase in helicase activity would be expected to reduce telomere extension, and we did not observe this. On the other hand, removal of stalled telomerase should free the enzyme, increasing extension activity at lower helicase concentrations, and gradually repressing extension as the concentration approaches levels where telomerase is unable to bind due to being overwhelmed by helicase removal activity. The latter scenario is most consistent with our telomerase assay results with both Pif1 (Fig. 2A, 4B) and Pif1+Hrq1 (Fig. 5B).

### Modulation of telomerase activity by tuning helicase concentration

*In vitro*, when telomerase binds to the 3ʹ end of telomeric ssDNA, it can extend the substrate via Type I processivity (Fig. 7). Subsequently, three things can happen: 1) telomerase can use Type II processivity to further extend the substrate; 2) telomerase can stall; or 3) telomerase can dissociate from the substrate and rebind at an internal position, leading to nuclease cleavage and Type I extension. Our data suggest that the concentrations of Hrq1 and Pif1 determine the outcome of these subsequent steps. At low concentrations of Hrq1 and Pif1, the helicases work together to stimulate telomerase activity (Fig. 5A-C). This is ostensibly due to inhibition or relief of telomerase stalling (non-productive), enabling processivity to occur (Fig. 7). However, at high Hrq1 and Pif1 concentrations, the two helicases become a potent telomerase inhibitor (Fig. 5A-C). This limits all types of processivity, likely through permanent eviction of telomerase from the ssDNA being patrolled by the helicases (Fig. 7). A similar process may also be used *in vivo*, where tuning the local concentration of Hrq1 and Pif1, not just within a nucleus but at an individual telomere, can be used to modulate telomerase activity. Low concentrations could stimulate telomerase activity, but high concentrations would inhibit telomere extension. As Pif1 is enriched at long telomeres *in vivo* (13), perhaps Hrq1 is as well. This would limit telomerase activity on long telomeres in favor of short ones, yielding a homeostatic telomere length. Although not depicted in Figure 7, our experiments with ATPase-null helicases also suggest that tuning the activity of Hrq1 and Pif1 can also modulate telomerase activity (Fig. 5, 6). *In vivo*, this could be accomplished by post-translational modifications of the helicases, and phosphorylation of Pif1 is already known to affect its activity at telomeres and DSBs (57,58).

**Figure 7.**
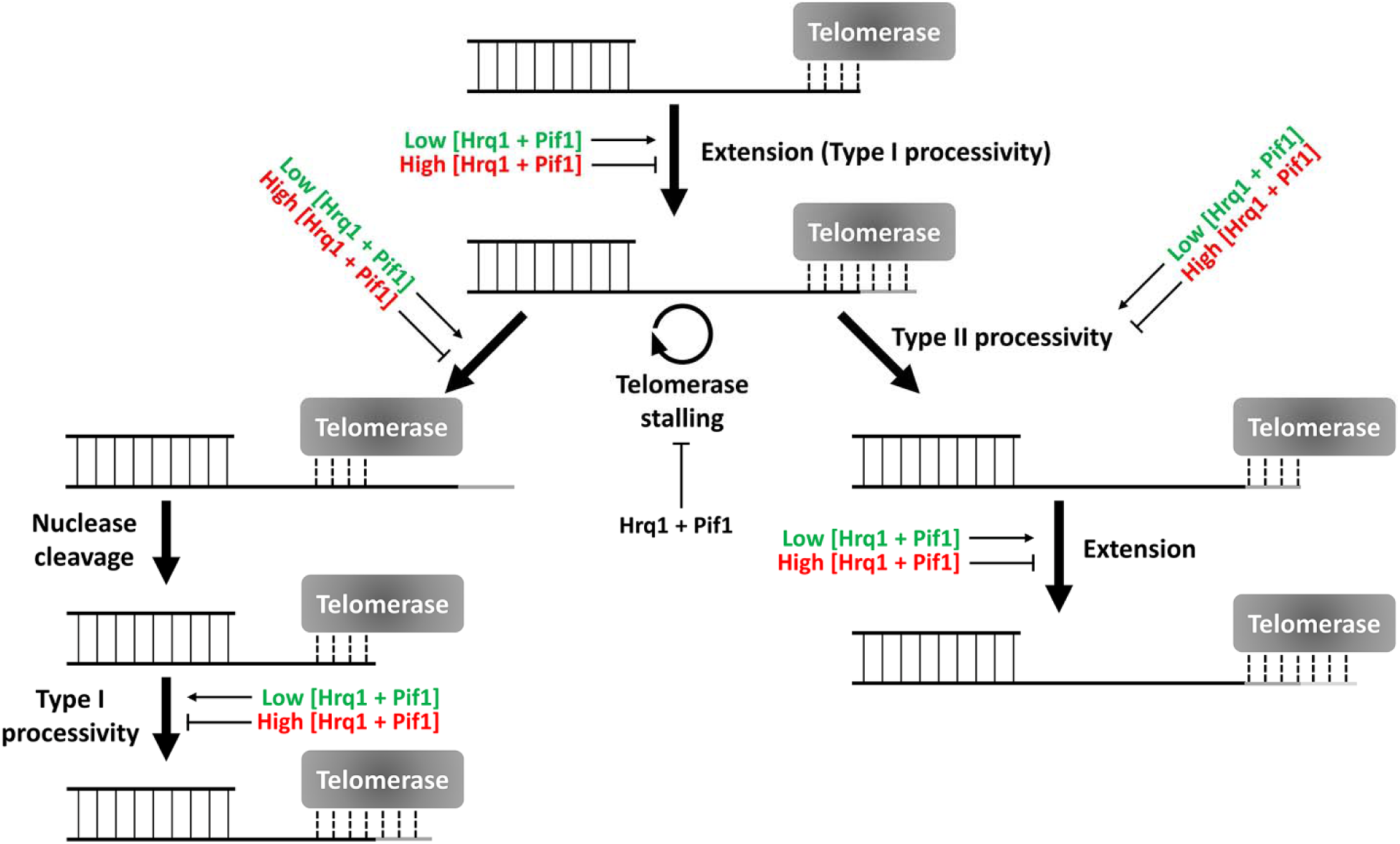
Model for how helicase concentration affects telomere length homeostasis. As stated in the text, telomerase can reiteratively extend telomeric ssDNA (Type I and II processivity), stall after extension, or dissociate and rebind internally to the telomeric ssDNA. A stall may occur in the latter case, which can lead to nuclease cleavage of the ssDNA, allowing telomerase to extend the substrate by Type I processivity again. Low concentrations of Hrq1 and Pif1 together stimulate telomerase extension *in vitro*, but high concentrations of both helicases are inhibitory. Both helicase, alone and in tandem, inhibit stalling.

Ultimately, the mechanism of telomere length homeostasis will remain a mystery until a completely reconstituted *in vitro* system is established. This will require the use of additional recombinant proteins such as the Ku70/80 heterodimer and the proteins and complexes that specifically bind double-stranded and ssDNA at telomeres (59,60), as well as the purified telomerase holoenzyme. Such investigations will shed light on the roles of these proteins in genome integrity, cellular aging, and disease.

## EXPERIMENTAL PROCEDURES

### Strains, Media, and Reagents

*S. cerevisiae* strain JBBY26, a derivative of BCY123 (*MATa can1 ade2 trp1 ura3-52 his3 leu2-3, 112 pep4::HIS3 prb1::LEU2 bar1::HISG, lys2::pGAL1/10-GAL4*) (61), harbors a dual over-expression plasmid for *TLC1* and *EST2. Escherichia coli* strain Rosetta 2 (DE3) pLysS (Novagen) was used for the over-expression of SUMO-tagged Pif1 and SUMO protease. Yeast cells were grown in SC-Ura drop out media for propagation and Est2/TLC1 overproduction. Rosetta cells were maintained on LB medium supplemented with 50 μg/mL kanamycin and 34 μg/mL chloramphenicol. Liquid cultures were grown in 2x YT medium for protein overproduction and supplemented with the same antibiotics. Radiolabeled ^32^P-[α]-TTP, ^32^P-[α]-ATP, and ^32^P-[γ]-ATP were purchased from PerkinElmer (Waltham, MA). All dNTPs were purchased from NEB (Ipswich, MA). Oligonucleotides were purchased from IDT (Coralville, IA), and the Tel15, Tel30, and Tel50 primers used for quantitative telomerase assays were PAGE purified. All primers used are listed in Table 1. Chemical reagents were purchased from Sigma-Aldrich (St. Louis, MO) or DOT Scientific (Burton, MI).

### Protein Purification and Enrichment

Plasmid pJBBY26, the media used for overproduction of Est2/*TLC1* in yeast, and the telomerase enrichment protocol have been previously described (23). Plasmids pSUMO-Pif1, pSUMO-Pif1-K264A, and pUlp1 were used for the overproduction of SUMO-tagged Pif1, the K264A mutant, and SUMO protease, respectively. Plasmid pSUMO-Pif1 was a gift from Kevin Raney. This plasmid was used as the template for standard site-directed mutagenesis to create pSUMO-Pif1-K264A. All three proteins were over-expressed in Rosetta cells grown overnight at 18°C with 0.2 mM IPTG induction. Cultures were centrifuged at 7000 × g, and cell pellets were resuspended in lysis buffer (50 mM sodium phosphate buffer (pH 7.5), 300 mM NaCl, 5 mM β-mercaptoethanol, 10% glycerol (w/v), protease inhibitor cocktail, and 0.5 mg/mL lysozyme). Resuspended cells were lysed in a cell cracker, and lysates were subjected to ultracentrifugation at 150,000 × g for 1 h. Supernatants were partially purified on a 1-mL Talon HiTrap column and eluted with 200 mM imidazole. Eluates enriched with helicase were pooled, dialyzed to remove imidazole, and digested for 2 h with the SUMO protease Ulp1. The Ulp1-digested samples were run over a Talon HiTrap column, and the flow through containing the untagged helicase, was collected. Uncleaved SUMO-Pif1, cleaved SUMO tag, and SUMO-Ulp1 remained bound to the column and were observed in imidazole-eluted fractions. Flow through fractions with helicase were diluted 1:1 into IEX buffer (50 mM sodium phosphate buffer (pH 6.8), 10% glycerol (w/v), 0.5 mM EDTA (pH 8.0), and 4 mM β-mercaptoethanol) to reduce the salt concentration to 150 mM. Samples were polished on a Resource S column and eluted in IEX buffer supplemented with NaCl using a gradient from 150 mM to 1 M. Samples were concentrated with Amicon Ultra-4 centrifugal filters with a 30K cutoff and stored at − 80°C in storage buffer (25 mM HEPES (pH 7.5), 150 mM NaCl, 2 mM β-mercaptoethanol, 0.1 mM EDTA (pH 8.0), and 30% glycerol (w/v)). Hrq1 and Hrq1-K318A were overproduced in insect cells and purified as reported (15,62). All helicase preparations were tested for ATPase activity and the absence of contaminating nuclease activity before use in assays.

### Helicase Assays

Fork substrates for helicase assays were constructed by incubating two partially complementary oligonucleotides (both at 1 μM) overnight at 37°C for annealing. These substrates included DNA/DNA (oligonucleotides MB1169 and MB1170, Table 1) and DNA/RNA duplexes with ssDNA as the loading strand (oligonucleotides MB1360r and MB1170). RNase inhibitors (NEB) were used during the preparation of RNA-containing substrates. All reagents were prepared with DEPC-treated water. Helicase reactions were performed at 30°C for 30 min in 1 x binding buffer supplemented with 5 mM ATP. Reactions were stopped by mixing with 5 x dye-free load buffer and placed on ice. Labeled fork substrates were added to a final concentration of 0.1 nM. Helicase reaction products were separated on native 8% 19:1 acrylamide:bis-acrylamide gels supplemented with 10 mM MgOAc and 5% glycerol. The Tris-Borate-EDTA running buffer (45 mM Tris-borate, 1 mM EDTA, pH 8.0) was supplemented with 2.5 mM MgOAc. Gels were prepared as described above.

### Telomerase Assays

Telomerase-enriched extracts were prepared by DEAE fractionation of clarified lysates (11,23). Briefly, Est2/TLC1 were overproduced by galactose induction, and cell pellets were prepared by centrifugation at 3000 × g for 10 min at 4°C. The pellets were resuspended in 1 mL of L buffer (40 mM Tris-HCl (pH 8.0), 500 mM NaOAc, 1.1 mM MgCl2, 0.1 mM EDTA, 0.1% (w/v) Triton-X100, 0.2% (w/v) NP-40 substitute, 10% (w/v) glycerol, and 0.1 mM PMSF) per mL of cell paste and flash frozen in liquid nitrogen. Frozen pellets were lysed by grinding with dry ice, slowly thawed at 4°C, and clarified by centrifugation at 10,000 × g. Clarified lysates were incubated with DEAE Sepharose Fast Flow (GE Healthcare) that was pre-equilibrated with TMG500 buffer (10 mM Tris-HCl (pH 8.0), 500 mM NaOAc, 2.2 mM MgCl_2_, 0.1 mM EDTA, 0.1% (w/v) Triton-X100, 0.2% (w/v) NP-40 substitute, 10% (w/v) glycerol, and 0.1 mM PMSF). After 30 min at 4°C, the DEAE resin was centrifuged at 800 × g for 1 min and washed three times with TMG500 buffer. Telomerase was eluted by a high salt wash with TMG900 (TMG supplemented with 900 mM sodium acetate). The eluate was concentrated an Amicon Ultra-4 centrifugal filter with a 30K cutoff to approximately 4 mL. Partially concentrated eluate was desalted using a 4-mL Zeba spin column (Thermo Scientific). Following desalting, samples were further concentrated to approximately 100 μL/L of starting culture. Samples were diluted 1:1 into glycerol, flash-frozen in liquid nitrogen, and stored at −80°C. Each telomerase preparation was titrated to standardize activity levels before use in experiments. Control experiments confirmed that all telomerase extension products were RNaseA sensitive (data not shown). Additional controls with only [^32^P]-α-TTP added had a predominant +1 nt extension product as expected for telomerase elongation from Tel15 (data not shown). Primer concentration was also titrated for activity. Based on these experiments, we prepared all telomerase assays with 1 μM primer, which was optimal for activity and ensured that helicase concentrations were sub-stoichiometric with respect to added primer.

Telomerase reactions were performed in 10 μL of 1 x telomerase reaction buffer (20 mM Tris-HCl (pH 8.0), 20 mM NaCl, 5 mM MgCl_2_, 1 mM DTT, 1 mM spermidine, 1 μL of 10 μM telomeric primer, 50 μM dGTP, 5 mM TTP, 1 μL of α-^32^P-dTTP (10 μCi/μL), RNase inhibitor (1U/μL) and 1 to 3 μL of telomerase-enriched extract) and incubated at 30°C for 45 min. In reactions with two primers, each was used at 0.5 μM. Reactions were stopped by adding 25 volumes of stop buffer (20 mM Tris-HCl (pH 8.0), 1 mM EDTA, 0.5% (w/v) SDS, 250 μg/mL proteinase K, and approximately 1000 cpm of ^32^γ-labeled loading control oligonucleotide) and incubating at 30°C for 45 min. Samples were extracted twice with phenol/chloroform (pH 7.0) and precipitated by addition of 1 volume of 4 M (NH_4_)_2_OAc and 2.5 volumes of ice-cold 100% ethanol. Tubes were chilled at −80°C for ≥30 min and centrifuged at 21,130 x g to precipitate reaction products. Pellets were resuspended in 6μL of 95% (v/v) formamide load buffer, heated to 95°C for 5 min, and reaction products were separated on 16% 19:1 acrylamide:bis-acrylamide gels containing 6 M urea. Gels were run at 2500 V for 120 min, dried, and imaged and quantified using a Typhoon 9500 scanner with ImageQuant software. For combined helicase and telomerase assays, the indicated helicase was diluted into 1 × Hrq1 storage buffer (25 mM Na-HEPES (pH 8.0), 30% glycerol, 300 mM NaOAc (pH 7.6), 25 mM NaCl, 5 mM MgOAc, 1 mM DTT, and 0.1% Tween-20) so that a 10-fold dilution into reactions achieved the desired helicase concentration. Control reactions lacking added helicase received 1 μL of 1 × Hrq1 storage buffer without added protein.

Total activity was measured by densitometry for each band on a gel using ImageQuant. The sum of the measured values for each band in a lane is reported as the total activity. Bands were corrected for the number of dT residues (*i.e*., the amount of α-^32^P-dTTP incorporation) and normalized to a loading control to generate corrected pixel values. The distribution of label was determined by adding the corrected pixels for each individual band to obtain total cumulative activity. Each individual band was then divided by the total activity to determine the percent of label in each band, referred to as label distribution. Telomerase extensions T51-58 from experiments with Tel50 were quantified as a group due to low resolution of the bands. Processivity was calculated by submitting the corrected pixel data to the following calculation: (T_I_+1…T_n_)/(T_I_+…T_n_), where T_I_ is the corrected pixels of lowest molecular weight telomerase extension band, and T_n_ is the highest molecular weight band in a series.

### Electrophoresis Mobility Assays (EMSAs)

Substrates for EMSAs were prepared by end labeling oligonucleotides Tel15, Tel30, Tel50, or MB1170 (Table 1). All “Tel” oligonucleotides contained the *S. cerevisiae* telomere repeat sequence TG_1-3_. Oligonucleotides were labeled with T4 polynucleotide kinase and γ-^32^P-ATP under standard conditions. Labeled oligonucleotides were separated from unincorporated label using G50 micro-columns (GE Healthcare). Binding reactions were performed in 1 x binding buffer (25 mM HEPES (pH 8.0), 5% glycerol (w/v), 50 mM NaOAc, 150 μM NaCl, 7.5 mM MgCl_2_, and 0.01% Tween-20 (w/v)). Radiolabeled substrates were boiled, placed on ice, and added to binding reactions to a final concentration 0.2 nM. When present, ATP was added to binding reactions to a final concentration of 4 mM. Binding reactions were incubated at 30°C for 30 min and mixed with 5 x dye-free loading buffer (50 mM Tris (pH 8.0) and 25% glycerol (w/v)). The reactions were separated on native 4% 37.5:1 acrylamide:bis-acrylamide gels in 1 x Tris-glycine running buffer (25 mM Tris (pH 8.0) and 185 mM glycine, pH 8.8). Gels were run at 100 V for 30-45 min, dried, and imaged and quantified using a Typhoon 9500 scanner with ImageQuant software. Binding reactions with Tel50 primer were separated on 0.5% agarose gels in a 1x TGE buffer (25 mM Tris and 185 mM glycine, pH 8.76) supplemented with 1 mM MgCl_2_. All data were plotted and, where appropriate, fit with curves using GraphPad software.

### Statistical analyses

All data were analyzed and graphed using GraphPad Prism 6 software. The reported values are averages of ≥3 independent experiments, and the error bars are the standard deviation. *P*-values were calculated as described in the figure legends, and we defined statistical significance as *p*<0.01.

## Acknowledgements

We thank members of the Bochman lab and Julia van Kessel for thoughtful comments on this manuscript.

## Conflict of interest

The authors declare that they have no conflicts of interest with the contents of this article.

## FOOTNOTES

The work in this paper was supported by funds from the College of Arts And Sciences, Indiana University, a Collaboration in Translational Research Pilot Grant from the Indiana Clinical and Translational Sciences Institute, and the American Cancer Society (RSG-16-180-01-DMC).

The abbreviations used as: ssDNA, single-stranded DNA; DSBs, double-strand breaks; GCR, gross-chromosomal rearrangement; STEX, single telomere extension

## REFERENCES

1. Dey, A., and Chakrabarti, K. (2018) Current Perspectives of Telomerase Structure and Function in Eukaryotes with Emerging Views on Telomerase in Human Parasites. Int J Mol Sci 19

2. Watson, J. D. (1972) Origin of concatameric T7 DNA. Nature New Biology 239, 197–201

3. Pardue, M. L., Rashkova, S., Casacuberta, E., DeBaryshe, P. G., George, J. A., and Traverse, K. L. (2005) Two retrotransposons maintain telomeres in Drosophila. Chromosome Res 13, 443–453

4. Armstrong, C. A., and Tomita, K. (2017) Fundamental mechanisms of telomerase action in yeasts and mammals: understanding telomeres and telomerase in cancer cells. Open Biol 7

5. Leao, R., Apolonio, J. D., Lee, D., Figueiredo, A., Tabori, U., and Castelo-Branco, P. (2018) Mechanisms of human telomerase reverse transcriptase (hTERT) regulation: clinical impacts in cancer. J Biomed Sci 25, 22

6. Eitsuka, T., Nakagawa, K., Kato, S., Ito, J., Otoki, Y., Takasu, S., Shimizu, N., Takahashi, T., and Miyazawa, T. (2018) Modulation of Telomerase Activity in Cancer Cells by Dietary Compounds: A Review. Int J Mol Sci 19

7. Li, H., Zhao, L., Yang, Z., Funder, J. W., and Liu, J. P. (1998) Telomerase is controlled by protein kinase Calpha in human breast cancer cells. J Biol Chem 273, 33436–33442

8. Kim, J. H., Park, S. M., Kang, M. R., Oh, S. Y., Lee, T. H., Muller, M. T., and Chung, I. K. (2005) Ubiquitin ligase MKRN1 modulates telomere length homeostasis through a proteolysis of hTERT. Genes Dev 19, 776–781

9. MacNeil, D. E., Bensoussan, H. J., and Autexier, C. (2016) Telomerase Regulation from Beginning to the End. Genes (Basel) 7

10. Schulz, V. P., and Zakian, V. A. (1994) The *Saccharomyces PIF1* DNA helicase inhibits telomere elongation and *de novo* telomere formation. Cell 76, 145–155

11. Boule, J., Vega, L., and Zakian, V. (2005) The Yeast Pif1p helicase removes telomerase from DNA. Nature 438, 57–61

12. Boule, J. B., and Zakian, V. A. (2007) The yeast Pif1p DNA helicase preferentially unwinds RNA DNA substrates. Nucleic Acids Res 35, 5809–5818

13. Li, J. R., Yu, T. Y., Chien, I. C., Lu, C. Y., Lin, J. J., and Li, H. W. (2014) Pif1 regulates telomere length by preferentially removing telomerase from long telomere ends. Nucleic Acids Res 42, 8527–8536

14. Bochman, M. L., Paeschke, K., Chan, A., and Zakian, V. A. (2014) Hrq1, a Homolog of the Human RecQ4 Helicase, Acts Catalytically and Structurally to Promote Genome Integrity. Cell reports 6, 346–356

15. Rogers, C. M., Wang, J. C., Noguchi, H., Imasaki, T., Takagi, Y., and Bochman, M. L. (2017) Yeast Hrq1 shares structural and functional homology with the disease-linked human RecQ4 helicase. Nucleic Acids Res 45, 5217–5230

16. Myung, K., Chen, C., and Kolodner, R. D. (2001) Multiple pathways cooperate in the suppression of genome instability in *Saccharomyces cerevisiae*. Nature 411, 1073–1076

17. Zhou, J.-Q., Monson, E. M., Teng, S.-C., Schulz, V. P., and Zakian, V. A. (2000) The Pif1p helicase, a catalytic inhibitor of telomerase lengthening of yeast telomeres. Science 289, 771–774

18. Ferrarelli, L. K., Popuri, V., Ghosh, A. K., Tadokoro, T., Canugovi, C., Hsu, J. K., Croteau, D. L., and Bohr, V. A. (2013) The RECQL4 protein, deficient in Rothmund-Thomson syndrome is active on telomeric D-loops containing DNA metabolism blocking lesions. DNA Repair (Amst)

19. Ghosh, A. K., Rossi, M. L., Singh, D. K., Dunn, C., Ramamoorthy, M., Croteau, D. L., Liu, Y., and Bohr, V. A. (2012) RECQL4, the protein mutated in Rothmund-Thomson syndrome, functions in telomere maintenance. J Biol Chem 287, 196–209

20. Dietschy, T., Shevelev, I., and Stagljar, I. (2007) The molecular role of the Rothmund-Thomson-, RAPADILINO- and Baller-Gerold-gene product, RECQL4: recent progress. Cell Mol Life Sci 64, 796–802

21. Cohn, M., and Blackburn, E. H. (1995) Telomerase in yeast. Science 269, 396–400

22. Zhou, R., Zhang, J., Bochman, M. L., Zakian, V. A., and Ha, T. (2014) Periodic DNA patrolling underlies diverse functions of Pif1 on R-loops and G-rich DNA. eLife 3, e02190

23. Boule, J. B., and Zakian, V. A. (2010) Characterization of the Helicase Activity and Anti-Telomerase Properties of Yeast Pif1p *in vitro*, in Helicases Methods and Protocols, Methods in Molecular Biology Series, Humana Press, ed. pp 359–376

24. Srivatsan, A., Putnam, C. D., and Kolodner, R. D. (2018) Analyzing Genome Rearrangements in Saccharomyces cerevisiae. Methods Mol Biol 1672, 43–61

25. Peng, Y., Mian, I. S., and Lue, N. F. (2001) Analysis of telomerase processivity: mechanistic similarity to HIV-1 reverse transcriptase and role in telomere maintenance. Mol. Cell 7, 1201–1211.

26. Bosoy, D., and Lue, N. F. (2004) Yeast telomerase is capable of limited repeat addition processivity. Nucleic Acids Res 32, 93–101

27. Eugster, A., Lanzuolo, C., Bonneton, M., Luciano, P., Pollice, A., Pulitzer, J. F., Stegberg, E., Berthiau, A. S., Forstemann, K., Corda, Y., Lingner, J., Geli, V., and Gilson, E. (2006) The finger subdomain of yeast telomerase cooperates with Pif1p to limit telomere elongation. Nat Struct Mol Biol 13, 734–739

28. Tzfati, Y., Fulton, T. B., Roy, J., and Blackburn, E. H. (2000) Template boundary in a yeast telomerase specified by RNA structure. Science 288, 863–867

29. Forstemann, K., and Lingner, J. (2001) Molecular basis for telomere repeat divergence in budding yeast. Mol Cell Biol 21, 7277–7286

30. Melek, M., and Shippen, D. E. (1996) Chromosome healing: spontaneous and programmed de novo telomere formation by telomerase. Bioessays 18, 301–308

31. Niu, H., Xia, J., and Lue, N. F. (2000) Characterization of the interaction between the nuclease and reverse transcriptase activity of the yeast telomerase complex. Mol Cell Biol 20, 6806–6815

32. Collins, K., and Greider, C. W. (1993) *Tetrahymena* telomerase catalyzes nucleolytic cleavage and nonprocessive elongation. Genes Dev 7, 1364–1376

33. Oulton, R., and Harrington, L. (2004) A human telomerase-associated nuclease. Mol Biol Cell 15, 3244–3256

34. Collins, K., and Gandhi, L. (1998) The reverse transcriptase component of the Tetrahymena telomerase ribonucleoprotein complex. Proc Natl Acad Sci U S A 95, 8485–8490

35. Weinrich, S. L., Pruzan, R., Ma, L., Ouellette, M., Tesmer, V. M., Holt, S. E., Bodnar, A. G., Lichtsteiner, S., Kim, N. W., Trager, J. B., Taylor, R. D., Carlos, R., Andrews, W. H., Wright, W. E., Shay, J. W., Harley, C. B., and Morin, G. B. (1997) Reconstitution of human telomerase with the template RNA component hTR and the catalytic protein subunit hTRT. Nat Genet 17, 498–502

36. Phillips, J. A., Chan, A., Paeschke, K., and Zakian, V. A. (2015) The Pif1 helicase, a negative regulator of telomerase, acts preferentially at long telomeres. PLoS Genet 11, e1005186

37. Barranco-Medina, S., and Galletto, R. (2010) DNA binding induces dimerization of Saccharomyces cerevisiae Pif1. Biochemistry 49, 8445–8454

38. Bochman, M. L., Sabouri, N., and Zakian, V. A. (2010) Unwinding the functions of the Pif1 family helicases. DNA Repair (Amst) 9, 237–249

39. Chib, S., Byrd, A. K., and Raney, K. D. (2016) Yeast Helicase Pif1 Unwinds RNA:DNA Hybrids with Higher Processivity than DNA:DNA Duplexes. J Biol Chem 291, 5889–5901

40. Greider, C. W. (2016) Regulating telomere length from the inside out: the replication fork model. Genes Dev 30, 1483–1491

41. Strecker, J., Stinus, S., Caballero, M. P., Szilard, R. K., Chang, M., and Durocher, D. (2017) A sharp Pif1-dependent threshold separates DNA double-strand breaks from critically short telomeres. eLife 6

42. Larrivee, M., LeBel, C., and Wellinger, R. J. (2004) The generation of proper constitutive G-tails on yeast telomeres is dependent on the MRX complex. Genes Dev. 18, 1391–1396

43. Chung, W. H., Zhu, Z., Papusha, A., Malkova, A., and Ira, G. (2010) Defective resection at DNA double-strand breaks leads to de novo telomere formation and enhances gene targeting. PLoS Genet 6, e1000948

44. Hardy, J., Churikov, D., Geli, V., and Simon, M. N. (2014) Sgs1 and Sae2 promote telomere replication by limiting accumulation of ssDNA. Nature communications 5, 5004

45. Wellinger, R. J., and Zakian, V. A. (2012) Everything you ever wanted to know about Saccharomyces cerevisiae telomeres: beginning to end. Genetics 191, 1073–1105

46. Zhang, D. H., Zhou, B., Huang, Y., Xu, L. X., and Zhou, J. Q. (2006) The human Pif1 helicase, a potential *Escherichia coli* RecD homologue, inhibits telomerase activity. Nucl Acids Res 34, 1393–1404

47. Barefield, C., and Karlseder, J. (2012) The BLM helicase contributes to telomere maintenance through processing of late-replicating intermediate structures. Nucleic Acids Res 40, 7358–7367

48. Drosopoulos, W. C., Kosiyatrakul, S. T., and Schildkraut, C. L. (2015) BLM helicase facilitates telomere replication during leading strand synthesis of telomeres. J Cell Biol 210, 191–208

49. Damerla, R. R., Knickelbein, K. E., Strutt, S., Liu, F. J., Wang, H., and Opresko, P. L. (2012) Werner syndrome protein suppresses the formation of large deletions during the replication of human telomeric sequences. Cell Cycle 11, 3036–3044

50. Edwards, D. N., Machwe, A., Chen, L., Bohr, V. A., and Orren, D. K. (2015) The DNA structure and sequence preferences of WRN underlie its function in telomeric recombination events. Nature communications 6, 8331

51. Opresko, P. L. (2008) Telomere ResQue and preservation--roles for the Werner syndrome protein and other RecQ helicases. Mech Ageing Dev 129, 79–90

52. Azam, M., Lee, J. Y., Abraham, V., Chanoux, R., Schoenly, K. A., and Johnson, F. B. (2006) Evidence that the S. cerevisiae Sgs1 protein facilitates recombinational repair of telomeres during senescence. Nucleic Acids Res 34, 506–516

53. Huang, P., Pryde, F., Lester, D., Maddison, R., Borts, R., Hickson, I., and Louis, E. (2001) SGS1 is required for telomere elongation in the absence of telomerase. Curr. Biol. 11, 125–129

54. Johnson, F., Marciniak, R., McVey, M., Stewart, S., Hahn, W., and Guarente, L. (2001) The *Saccharomyces cerevisiae* WRN homolog Sgs1p participates in telomere maintenance in cells lacking telomerase. EMBO J. 20, 905–913

55. Lydeard, J. R., Lipkin-Moore, Z., Jain, S., Eapen, V. V., and Haber, J. E. (2010) Sgs1 and exo1 redundantly inhibit break-induced replication and de novo telomere addition at broken chromosome ends. PLoS Genet 6, e1000973

56. Pickett, H. A., and Reddel, R. R. (2012) The role of telomere trimming in normal telomere length dynamics. Cell Cycle 11, 1309–1315

57. Makovets, S., and Blackburn, E. H. (2009) DNA damage signalling prevents deleterious telomere addition at DNA breaks. Nat Cell Biol 11, 1383–1386

58. Vasianovich, Y., Harrington, L. A., and Makovets, S. (2014) Break-induced replication requires DNA damage-induced phosphorylation of Pif1 and leads to telomere lengthening. PLoS Genet 10, e1004679

59. Lewis, K. A., and Wuttke, D. S. (2012) Telomerase and telomere-associated proteins: structural insights into mechanism and evolution. Structure 20, 28–39

60. Nandakumar, J., and Cech, T. R. (2013) Finding the end: recruitment of telomerase to telomeres. Nat Rev Mol Cell Biol 14, 69–82

61. Zhao, S., Douglas, N. W., Heine, M. J., Williams, G. M., Winther-Larsen, H. C., and Meaden, P. G. (1994) The STL1 gene of Saccharomyces cerevisiae is predicted to encode a sugar transporter-like protein. Gene 146, 215–219

62. Rogers, C. M., and Bochman, M. L. (2017) Saccharomyces cerevisiae Hrq1 helicase activity is affected by the sequence but not the length of single-stranded DNA. Biochem Biophys Res Commun 486, 1116–1121

